# Genomic and Evolutionary Determinants of Two-hit Frequencies in Tumor Suppressor Genes

**DOI:** 10.64898/2026.02.17.706175

**Authors:** Nivedita Mukherjee, Radhakrishnan Sabarinathan

## Abstract

While biallelic or “two-hit” inactivation is a central organizing principle for tumor suppressor genes (TSGs), large-scale cancer genomic data reveal substantial heterogeneity in the frequency of such events across genes. This variability reflects diverse selective constraints on allelic disruption, whose biological determinants remain incompletely characterized. Here, we present a comprehensive, allele-specific analysis of TSG two-hit alterations across ∼9,000 tumors from The Cancer Genome Atlas, focusing on loss of heterozygosity (LOH) arising from the co-occurrence of point mutation and deletion. We show that two-hit frequencies vary widely across TSGs and scale with the functional impact and selection strength of point mutations. Integrating mutation position with zygosity reveals distinct patterns consistent with dominant versus recessive modes of action, enabling a zygosity-informed framework for variant interpretation. We further demonstrate that chromosomal context strongly shapes two-hit frequencies, reflecting aneuploidy biases across chromosome arms and selective trade-offs imposed by neighboring loci, including the co-deletion of synergistic TSGs. Extending the LOH analysis beyond diploid tumors, we find that equivalent “all-hit” frequencies are largely preserved in polyploid cancers following whole-genome doubling, consistent with early acquisition of LOH during clonal evolution. Collectively, our results uncover multiple determinants of adherence to the two-hit model, providing new insight into long-standing heterogeneity in TSG behavior and its potential clinical relevance.

## Introduction

Cancer development is an evolutionary process in which somatic cells accumulate genetic alterations and undergo clonal expansion or contraction shaped by natural selection^1^. Accordingly, genetic alterations observed in cancer genomes broadly fall into two classes: ‘drivers’, which confer a fitness advantage to the cell by perturbing cellular processes underlying the ‘hallmarks of cancer’, and ‘passengers’, which are selectively neutral and evolve largely through genetic drift^2,3^. Driver alterations are often identified based on predicted functional impact, and recurrence patterns across patient cohorts that reflect signals of positive selection, and form key targets in many cancer screening and precision medicine strategies^4,5^. Genes harboring these driver alterations include oncogenes (ONGs), which promote cell proliferation through activating mutations, and tumor suppressor genes (TSGs), which normally restrain proliferation and are inactivated in cancer. Unlike ONGs, which act dominantly and generally present as monoallelic alterations, malignant transformation driven by TSGs require alteration of both alleles to achieve complete inactivation^6^. This principle, formalized as Knudson’s two-hit hypothesis, has long been a cornerstone of cancer genetics^6^.

However, large-scale cancer genomic studies have revealed striking variability in the frequency of biallelic alterations across TSGs^7^. While some TSGs are almost invariably inactivated through two-hits, others frequently exhibit only a single hit, with one allele remaining intact. High two-hit frequencies likely reflect strong positive selection for complete gene inactivation, whereas lower frequencies may indicate negative selection or simply the lack of positive selection. For example, “one-hit” TSG drivers, such as those acting through haploinsufficiency, dominant-negative, or gain-of-function mechanisms, may not require a second hit, partially accounting for this variability^8^. Alternatively, these patterns may also arise from non-selective processes or genetic drift. However, despite the scale of available cancer genomic datasets, systematic studies surveying the extent of deviations from the two-hit hypothesis across TSGs and dissecting the factors underlying variability in two-hit frequencies remain limited^9,10^.

Towards this effort, we conducted a comprehensive, allele-specific analysis of TSG alterations using data from The Cancer Genome Atlas (TCGA) cohort, encompassing approximately 9,000 patients across 33 cancer types^11,12^. We focused on the most prevalent form of biallelic alteration—point mutation in one allele co-occurring with deletion of the other—resulting in an allelic configuration referred to as loss of heterozygosity (LOH). By characterizing the spectrum of alterations associated with LOH, we discovered that the two-hit frequencies scale with the functional impact and inferred selection strength of point mutations. We further illustrated how this relationship can be interpreted along with the mutation position to inform variant interpretation. Besides, we show that deletions contributing to LOH are rarely localized events and often comprise aneuploidies that affect neighboring loci. This makes two-hit frequencies strongly dependent on the chromosomal context of the gene, including selective advantage from the co-deletion of linked TSG clusters. Finally, since LOH is traditionally conceptualized within a diploid framework, prior studies have largely overlooked how LOH rates are affected by polyploidy following events such as whole-genome doubling (WGD). By tracing clonal dynamics in polyploid tumors, we find that TSG alterations occur early during evolution, thereby maintaining consistent LOH frequencies despite genome doubling. Collectively, our results illuminate the genomic factors and evolutionary forces shaping two-hit frequencies across TSGs in cancers.

## Results

### Co-occurring point mutations and deletions dominate TSG biallelic alterations

To characterize the landscape of biallelic alterations in TSGs, we analysed genetic alterations from 8,958 patients spanning 33 cancer types in the TCGA cohort (see Methods; **Fig. 1A**). We integrated allele-specific copy number profiles with point mutations, including somatic and pathogenic germline variants (substitutions and short indels), to infer the allelic configuration of each autosomal protein-coding gene. Based on prior knowledge, genes were classified into a high-confidence set of 132 TSGs, alongside negative control gene sets comprising 115 oncogenes (ONGs), 1,312 essential genes, 703 non-essential genes, and 12,536 unclassified genes (see Methods).

**Fig. 1:**
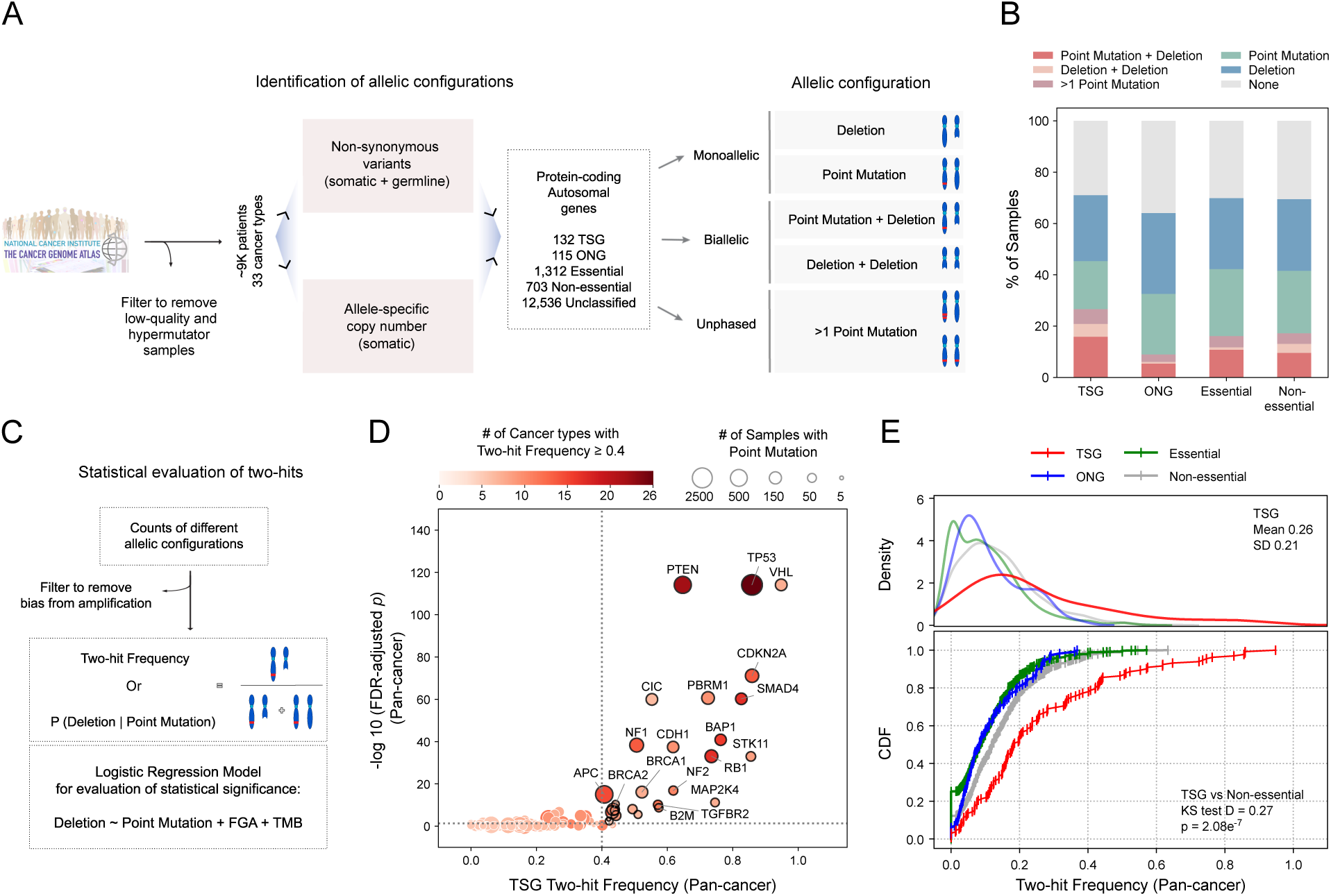
Allelic configurations and two-hit frequencies of TSGs. (A) Schematic of data processing steps to determine the allelic configurations or the alteration status of the alleles of protein-coding autosomal genes: tumor suppressor genes (TSG), oncogenes (ONG), essential genes, non-essential genes, and unclassified genes. (B) Prevalence of different allelic configurations across gene categories: biallelic alterations (Point Mutation + Deletion, Deletion + Deletion), monoallelic alterations (Point Mutation only, Deletion only), and unphased alterations (>1 Point Mutation). (C) Schematic illustrating the statistical evaluation of two-hit frequencies for each gene at pan-cancer or cancer type levels. Two-hit frequencies were defined as the conditional probability of observing a deletion given the presence of a point mutation: *P (Deletion | Point Mutation)*. Statistical significance was assessed using two-sided Wald tests derived from multivariable logistic regression adjusting for FGA (fraction of the genome altered) and TMB (tumor mutation burden). (D) Volcano plot showing pan-cancer two-hit frequencies of TSGs plotted against the negative log10 of FDR-adjusted (Benjamini-Hochberg method) Wald test *p*-values from the multivariate logistic regression model. Bubble size represents the number of samples harboring point mutations in each TSG, while bubble color indicates the number of cancer types in which the TSG exhibits a two-hit frequency ≥ 0.4. Horizontal and vertical dashed lines denote an FDR threshold of 0.05 and a two-hit frequency inflection point of 0.4, respectively. (E) Kernel Density Estimations or KDEs of pan-cancer two-hit frequencies across gene categories, with insets showing the mean and standard deviation of the TSG distribution (top). Cumulative Distribution Functions or CDFs of pan-cancer two-hit frequencies across gene categories, with insets displaying the Kolmogorov-Smirnov (KS) distance and *p*-value comparing TSGs and non-essential genes (bottom).

As expected, TSGs exhibited the highest incidence of biallelic alteration (21%), compared with ONGs (6%), essential genes (12%), and non-essential genes (13%) (*p* < 0.001, pairwise one-sample proportion tests; **Fig. 1B**). They also exhibited the lowest incidence of monoallelic alteration (44% vs 55%, 54% and 52% for TSGs vs ONGs, essential genes and non-essential genes, respectively; *p* < 0.001, pairwise one-sample proportion tests; **Fig. 1B**). Allelic configurations could not be resolved at loci harboring two or more point mutations and were therefore classified as unphased (6% for TSGs). Consistent with previous reports, biallelic alterations in TSGs predominantly resulted from the co-occurrence of a point mutation and deletion, leading to loss of heterozygosity (LOH), rather than from homozygous deletion of both alleles (16% vs 5%, respectively; **Fig. 1B**)^10^. This pattern was observed in 112 (85%) TSGs, with few known exceptions including FAS and CDKN2A (**Fig. S1A**)^7^. Hence, we focused on the most ubiquitous mechanism of biallelic alteration in TSGs, which is the co-occurrence of point mutation and deletion, hereafter referred to as “two-hit” events.

### Two-hit frequencies exhibit substantial heterogeneity across TSGs

We defined the two-hit frequency of each gene as the probability of observing a deletion in that gene, conditional on the presence of at least one non-synonymous point mutation on the remaining allele, i.e. P (Deletion | Point Mutation) (see Methods; **Fig. 1C**). However, such estimated frequencies may be confounded by sample-level variation in genome-wide point mutation and copy-number alteration levels. Consistent with this, cancer types with higher tumor mutation burden (TMB) exhibited lower TSG two-hit frequencies (Spearman’s ρ = −0.41, *p* = 0.024), whereas those with a greater fraction of the genome altered by copy-number changes (FGA) showed higher TSG two-hit frequencies (Spearman’s ρ = 0.46, *p* = 0.009; **Fig. S1B-C**). Hence, to account for this bias, we evaluated the statistical significance of mutation-deletion associations using a multivariate logistic regression model that adjusted for FGA and TMB (see Methods; **Fig. 1C**).

Pan-cancer two-hit frequencies of TSGs spanned a wide range, from 0 to 0.95, with 30 of the 132 genes exceeding 0.4 (**Fig. 1D, Fig. S2**). Notably, 27 of these 30 high-frequency TSGs (90%) showed statistically significant mutation-deletion associations (Wald test FDR < 0.05, multivariate logistic regression), making 0.4 the inflection point in the frequency versus FDR relationship. This group included genes with consistently high two-hit frequencies (≥ 0.4) across multiple cancer types (e.g. TP53, PTEN, SMAD4, RB1, BAP1, NF1, BRCA2, CDKN2A, STK11), as well as genes showing elevated frequencies only within specific cancer types (e.g. VHL, BRCA1, BRD7, MAP2K4, TSC1, CIC, AJUBA, MEN1, AXIN1) (**Fig. S2**). Of the remaining 102 TSGs, only 20 (20%) reached statistical significance (**Fig. 1D, Fig. S2**). Similarly, among the 256 TSG–cancer-type pairs with high two-hit frequencies (≥ 0.4), 77 (30%) were statistically significant, compared to only 8 of the remaining 534 pairs (1.5%) (**Fig. S2**). These results demonstrate concordance between observed two-hit frequencies and statistical significance after adjustment for FGA and TMB.

Overall, TSGs displayed higher and more variable pan-cancer two-hit frequencies (mean 0.26, SD 0.21) than all other gene categories, which exhibited lower and less dispersed distributions (means between 0.10 and 0.13, SDs between 0.09 and 0.10; TSG vs non-essential KS test *p* = 2.08e^-7^; **Fig. 1E**). A similar pattern was observed at the cancer type level as well (mean 0.30, SD 0.31 for TSGs; means between 0.10 and 0.13, SDs between 0.15 and 0.18 for all other gene categories; TSG vs non-essential KS test *p* = 7.66e^-25^; **Fig. S1D**). Hence, we conclude that TSGs acquire two-hits at markedly higher and more heterogenous rates than other gene categories. To uncover the genomic determinants of this variability, we dissected the point mutation and deletion patterns underlying elevated and reduced two-hit frequencies.

### Two-hit frequencies scale with the functional impact of TSG point mutations

Of the two classes of alterations contributing to TSG two-hits, deletions are expected to nearly always abolish gene function, whereas point mutations can have diverse functional outcomes. Consequently, the phenotypic impact and hence the associated selective advantage of a two-hit configuration may depend on the nature of the accompanying point mutation. To test this, we estimated selection coefficients separately for missense and truncating mutations in one-hit and two-hit configurations using the dNdScv algorithm (see Methods)^13^. This method infers selection during clonal evolution using the normalized ratio of non-synonymous to synonymous mutations (dN/dS), adjusted for variations in mutation rate across the genome. Assuming that synonymous mutations are selectively neutral and reflect background mutation rates, dN/dS > 1 indicates positive selection, dN/dS < 1 indicates negative selection, and dN/dS ≈ 1 indicates neutral evolution.

Two-hit–specific dN/dS values showed a strong positive correlation with the two-hit frequencies of TSGs, indicating that point mutations under stronger positive selection are associated with a higher probability of acquiring a second hit (ρ = 0.79 and *p* = 1.4e^-29^ for truncating, *ρ* = 0.75, p = 5.3e^-25^ for missense, Spearman’s correlation; **Fig. S3A-B**). Among the point mutations, truncating mutations were subject to substantially stronger positive selection than missense mutations, particularly for TSGs with high two-hit frequencies (**Fig. 2A**). Consistent with this pattern, 52 of 132 TSGs (∼39.4%) exhibited two-hit–specific dN/dS values significantly greater than 1 for truncating mutations, compared to only 16 TSGs (∼12%) for missense mutations. In fact, 15 TSGs, most with very low two-hit frequencies, showed evidence of negative selection for missense mutations in the two-hit configuration (**Fig. 2A**).

**Fig. 2:**
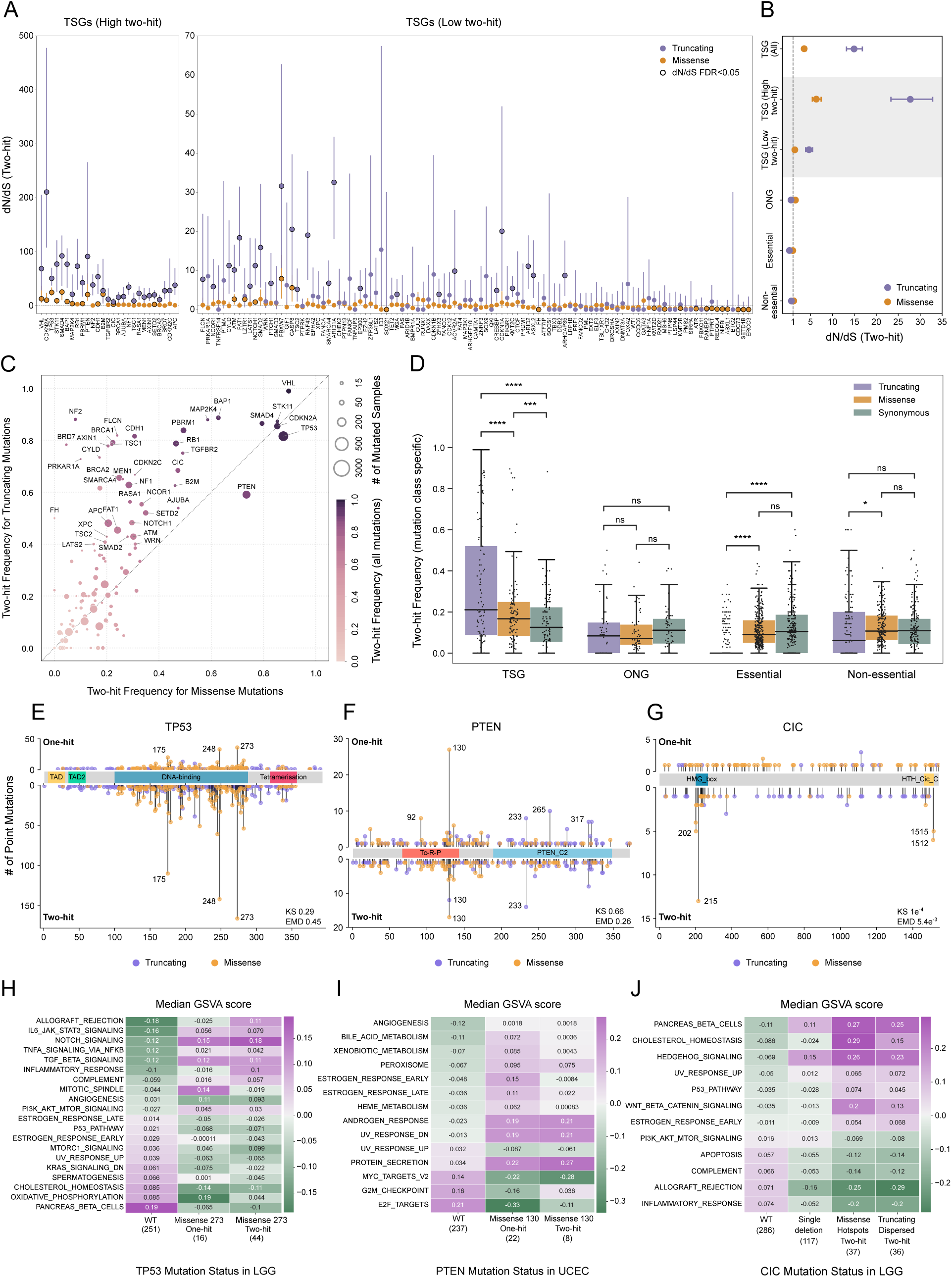
Functional impact of point mutations and two-hit frequencies. (A) Two-hit–specific dN/dS ratios for truncating (purple) and missense (orange) mutations in TSGs, ordered along the x-axis by decreasing pan-cancer two-hit frequency. TSGs with high two-hit frequencies (≥ 0.4, Wald test FDR < 0.05) are shown separately (left). Horizontal dashed line represents dN/dS value of 1. dN/dS ratios significantly different from 1 (FDR < 0.05, Likelihood ratio test) are outlined in black. (B) Global two-hit–specific dN/dS ratios for truncating (purple) and missense (orange) mutations in TSGs, ONGs, essential genes and non-essential genes. Global estimates calculated separately for TSGs with high and low two-hit frequencies are also shown. (C) Pan-cancer two-hit frequencies of TSGs computed separately for truncating and missense mutations. Bubble size reflects the number of samples with point mutations, and color reflects the two-hit frequency computed across all non-synonymous coding mutations. (D) Distributions of pan-cancer two-hit frequencies of different gene categories, computed separately for truncating, missense and synonymous mutations. Asterisks denote statistical significance based on paired Wilcoxon signed-rank tests: *p* > 0.05 (ns), ≤ 0.05 (*), ≤ 0.01 (**), ≤ 0.001 (***), ≤ 0.0001 (****). (E-G) Lollipop plots showing the distribution of point mutation positions along the linear polypeptide for one-hit and two-hit configurations in TP53 (E), PTEN (F), and CIC (G). X-axis represents amino acid position. Missense mutations are shown in yellow and truncating mutations in purple. Recurrent single-base hotspots (mutated in ≥ 5 patients) are annotated. Insets show p-values comparing one-hit and two-hit mutation position distributions for the mutation class of the most frequent hotspot in that gene, using Earth Mover’s Distance (EMD) and Kolmogorov–Smirnov (KS) tests. (H-I) Median GSVA enrichment scores for wild-type and mutant samples with TP53 R273 substitution in LGG (H) and PTEN R130 substitution in UCEC (I). Hallmark pathways significantly dysregulated between mutant and wild-type samples are shown. Mutant samples are divided into one-hit and two-hit configurations. Numbers in parentheses indicate the sample size for each comparison group. (J) Median GSVA enrichment scores for wild-type and CIC-mutant LGG samples. Hallmark pathways significantly dysregulated between samples with single-allele deletion and those with two-hit mutations are shown. Two-hit samples are divided into missense hotspot mutations and dispersed truncating mutations. Numbers in parentheses indicate the sample size for each comparison group.

A similar pattern was evident in the global dN/dS estimates (**Fig. 2B**). Across TSGs, the two–hit–specific selection coefficient for truncating mutations was 4.1-fold higher than that for missense mutations. This difference was even more pronounced for TSGs with high two-hit frequencies compared to those with lower frequencies (4.4-fold versus 3.3-fold). In contrast, in the control gene sets, truncating mutations were under negative selection, whereas missense mutations were either positively selected (ONGs) or neutrally evolving (essential and non-essential genes). In the one-hit configuration, global dN/dS estimates showed a similar relative trend, and 38 and 11 TSGs exhibited evidence of positive selection for truncating and missense mutations, respectively (**Fig. S3E-F**). However, one-hit–specific dN/ dS values showed no significant association with two-hit frequencies for either mutation class as expected (*p* = 0.56 for truncating, *p* = 0.85 for missense, Spearman’s correlation; **Fig. S3C-D**).

Next, to directly assess whether the functional consequence of point mutations is associated with variation in two-hit rates, we computed mutation-class–specific two-hit frequencies for truncating, missense, and synonymous mutations. Truncating mutations in TSGs exhibited higher two-hit frequencies than missense mutations, whereas synonymous mutations, as expected, showed the lowest frequencies (**Fig. 2C-D**, **Fig. S3G**). Notable exceptions included *PTEN*, *TP53*, and *CDKN2A*, for which two-hit frequencies were comparable between, or even higher for, missense than truncating mutations. We further stratified missense mutations by CADD PHRED score, a measure of predicted functional impact or deleteriousness relative to all possible human variants^14^. Missense mutations with high CADD scores displayed higher two-hit frequencies than those with low scores (**Fig. S3H**). As expected, these hierarchical patterns were absent in the control gene sets (**Fig. 2D**, **Fig. S3G-H**). For example, truncating mutations showed the lowest two-hit frequencies in essential genes, consistent with strong purifying selection (**Fig. 2D**, **Fig. S3G**).

The observed hierarchy between truncating and missense mutations likely reflects their distinct molecular consequences. Truncating mutations typically result in loss-of-function effects, which are recessive unless the gene is haploinsufficient, thereby conferring a selective advantage in the two-hit configuration. In contrast, missense mutations can have additional outcomes, including gain-of-function, dominant-negative, or neutral effects, which may not necessarily confer the same selective advantage^15^. Within both truncating and missense classes, two-hit frequencies scaled with the inferred strength of selection or deleteriousness of the mutations, as estimated by dN/dS values and CADD PHRED scores, respectively. Taken together, these analyses show that two-hit frequencies depend on the functional impact or mode of action of the accompanying point mutation, with higher-impact mutations or those likely to cause loss-of-function, being accompanied by deletion of the remaining allele more frequently than others.

### Spatial mutation patterns across one- and two-hit configurations inform point-mutation effects

In addition to mutation type, the position of a point mutation within a gene is a key determinant of its molecular consequence. To assess whether the probability of acquiring a two-hit configuration varies with mutation position, we compared the spatial distributions of point mutations along the linear polypeptide chain between one-hit and two-hit configurations (see Methods). Focusing on TSGs with missense or truncating hotspot mutations in either configuration, we identified multiple genes with significantly different spatial mutation patterns between the two configurations. Among these, several exhibited one-hit–specific hotspots, including *RNF43*, *CTCF*, *KMT2D*, *PIK3R1* and *FBXW7* (**Fig. S4A-E**). In contrast, two-hit–specific hotspots were observed in genes such as *CIC*, *VHL*, *SMAD4*, *B2M* and *BRCA2* (**Fig. 2G**, **S4F-I**). Given the previously observed association between point mutation consequences and two-hit frequencies, we reasoned that one-hit hotspot positions may reflect dominant modes of action, whereas two-hit hotspots may correspond to recessive loss-of-function effects. Consistent with this interpretation, TSGs identified with one-hit hotspots were also found to have evidence for gain-of-function, dominant-negative, or haploinsufficient effects, whereas mutations in those with two-hit hotspots are predominantly well-established loss-of-function events, with no comparable evidence for dominant effects^16–25^.

Among TSGs without significant difference in point mutation positions between one-hit and two-hit configurations, several displayed hotspot positions shared across both contexts. Examples include *TP53*, *PTEN, CDKN2A*, *APC* and *EP300*, most of which have also been implicated in dominant mechanisms (**Fig. 2E-F**, **Fig. S4J-L**)^26–29^. To further validate this interpretation, we examined transcriptomic signatures associated with the two hotspot mutations with the largest sample sizes: missense substitutions at TP53 R273 in LGG and PTEN R130 in UCEC (see Methods; **Fig. 2E-F**). Because deletion of the remaining allele should be phenotypically inconsequential when paired with a dominant mutation, we hypothesized that these hotspot mutations would exert similar functional effects in one-hit and two-hit configurations. LGG samples harboring TP53 R273 mutations showed significant dysregulation of 20 hallmark pathways compared with wild-type samples (FDR < 0.05, Mann–Whitney U test), yet none of these pathways differed significantly between one-hit and two-hit states (*p* < 0.05) (**Fig. 2H**). In both configurations, pathway changes occurred in the same direction relative to wild type. Similarly, in UCEC, 14 hallmark pathways were significantly dysregulated in samples with PTEN R130 mutations compared with wild type (FDR < 0.05), while none of them showed a significant difference between one-hit and two-hit configurations (*p* < 0.05) (**Fig. 2I**).

Upon further examination of TSGs with distinct spatial distributions of point mutations between the two-configurations, *CIC* stood out for its unique two-hit–specific hotspots^22,30^. In the two-hit configuration, missense mutations in *CIC* clustered at two distinct regions, whereas truncating mutations were distributed broadly along the polypeptide chain (**Fig. 2G**). This pattern, predominantly evident in LGG samples, suggests that two-hit events in *CIC* can arise either through position-agnostic truncating mutations or through missense mutations at specific sites that are sufficiently disruptive to cause loss-of-function. To test this, we compared transcriptomic signatures in LGG samples harboring two-hit *CIC* mutations with those containing a single-allele deletion of *CIC*, and identified 12 significantly dysregulated hallmark pathways (FDR < 0.05, Mann–Whitney U test). Only one of these affected pathways differed significantly between samples with the two-hit missense hotspot mutations and two-hit dispersed truncating mutations (cholesterol homeostasis pathway, *p* = 0.03, FDR = 0.3), indicating broadly similar transcriptomic effects across the two mutation classes (**Fig. 2J**). Notably, two-hit samples deviated from wild-type expression in the same direction as single-deletion samples, but with greater magnitude, consistent with a loss-of-function–like effect.

Collectively, these findings indicate that the probability of acquiring two hits is strongly influenced by the impact of the accompanying point mutation, which in turn is determined by its position within the polypeptide chain and its effect on the resultant protein. This relationship can be leveraged to help characterize point mutations, as illustrated by the case of *CIC*. However, it remains unclear why some dominantly acting point mutations appear exclusively as one-hit hotspots, whereas others are shared between one-hit and two-hit configurations. For example, although *TP53* has been reported to harbor dominant-negative and gain-of-function missense mutations^26,31^, which should not necessitate deletion of the other allele for phenotypic effect, we did not observe any hotspots that were strictly one-hit–specific. This observation is consistent with prior reports showing that dominant-negative *TP53* mutations do not alter the frequency of two-hit events^32^. While this could be due to dosage-sensitive effects, it also raises the possibility of additional determinants that modulate deletion patterns independent of the accompanying point mutation.

### Chromosomal properties constrain deletion lengths in TSG two-hits

Having characterized the association between point mutation types and two-hit frequencies, we next investigated the deletion patterns in TSG two-hit events. We determined deletion lengths and normalized them to chromosome arm length (see Methods). In cancer genomes, copy-number alterations typically occur as focal events, chromosome arm aneuploidies, and whole-chromosome aneuploidies, which appear in the normalized length distributions as peaks below 0.5, at 1, and at 2, respectively^33^. Analysis of TSG two-hit deletions revealed two groups of chromosomes: those enriched for focal deletions (chromosomes 1, 2, 4, 7, 11, 12, 16, and 20) and those enriched for arm aneuploidies (chromosomes 3, 5, 6, 8, 9, 10, 17, 18, and 19) (see Methods; **Fig. 3A**, **Fig. S5A**). Arm aneuploidies were largely confined to either the p or the q arm (**Fig. 3B**). Among acrocentric chromosomes, which have only q arms, chromosomes 14, 15, and 22 showed enrichment of arm-length over focal deletions, making arm aneuploidies the most common deletion mode in TSG two-hit events genome-wide (**Fig. 3B**). As expected, cancer-type–specific differences in aneuploidy patterns were observed, notably for arms 1p, 3p, 10q, and 13q (**Fig. S5B**)^34^. However, for each chromosome arm, the majority of cancer types retained the same arm-versus-focal enrichment pattern observed in the pan-cancer analysis.

**Fig. 3:**
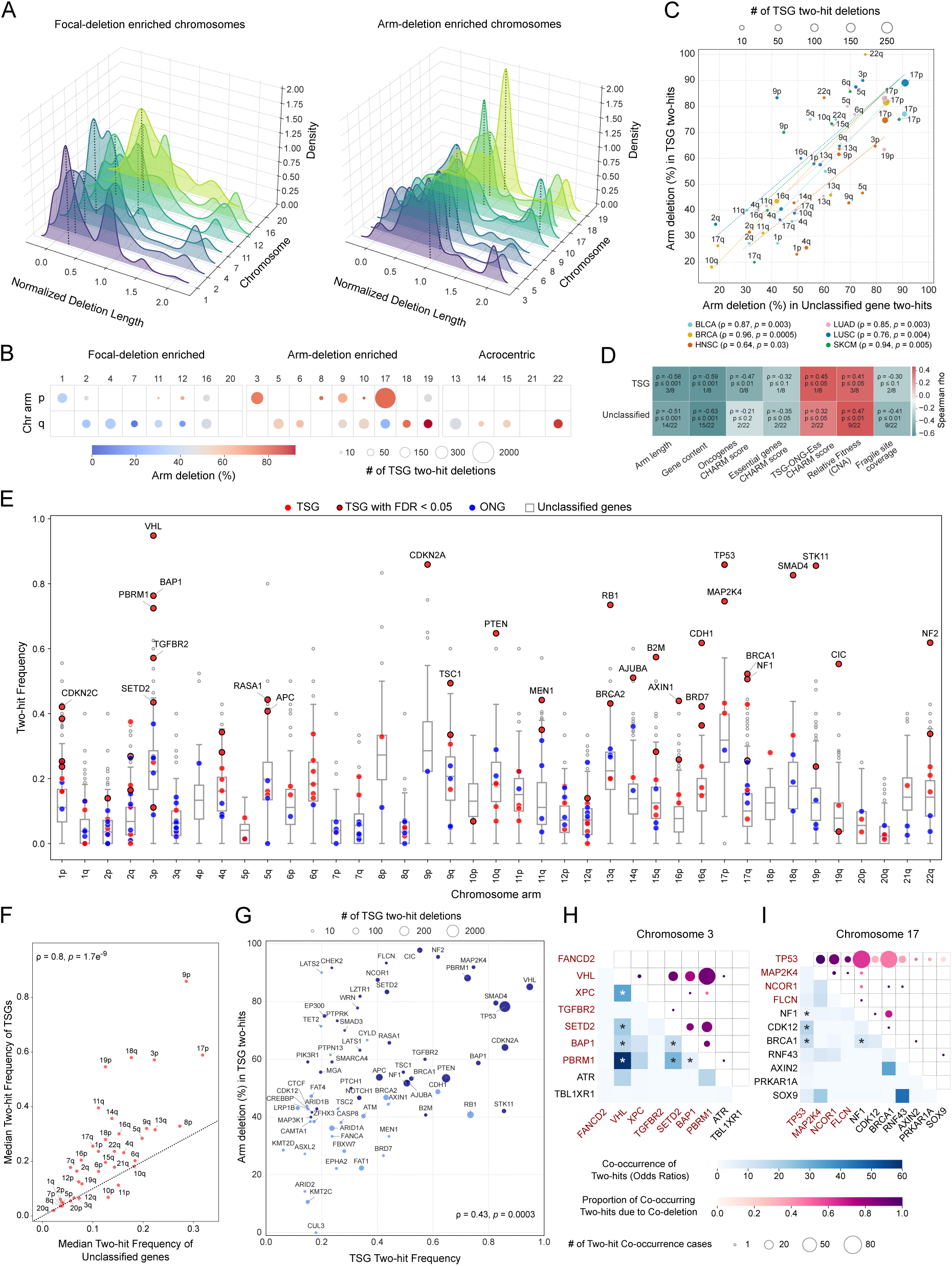
Chromosomal influence on two-hit frequencies and associated deletion patterns. (A) Kernel density estimation (KDE) plots showing normalized deletion-length distributions for TSG two-hit events across non-acrocentric chromosomes, grouped by enrichment for focal versus arm length deletions. Black dotted lines indicate the peak of each KDE curve. The area under each curve equals 1. (B) Prevalence of arm-length deletions in TSG two-hit events, defined as the percentage of deletions that are arm-length (≥ 70% of the arm) versus focal (≤ 50% of the arm), shown separately for p and q arms across arm-enriched, focal-enriched, and acrocentric chromosomes. Red bubbles indicate predominance of arm-length deletions, and blue bubbles indicate predominance of focal deletions. Bubble size reflects the total number of deletions (arm-length + focal). Only chromosome arms with ≥10 deletions are shown. (C) Comparison of arm-length deletion prevalence in two-hit events affecting TSGs versus unclassified genes, shown for each chromosome arm across cancer types. Bubble size denotes the total number of deletions, and bubble color indicates cancer type. Spearman’s correlation coefficients and p-values for each cancer type are provided in the legend. Of the 8 cancer types with at least five chromosome arms containing ≥ 10 deletions, the 6 with significant correlations (*p* < 0.05) are shown. Dashed lines represent linear fits for each cancer type; Pearson’s correlation *p*-values ranged from to 0.004 to 0.06, slopes ranged between 0.8 to 1. (D) Heatmap showing pan-cancer Spearman’s correlation coefficients between arm-length deletion prevalence (for TSGs and unclassified genes) and chromosome arm–intrinsic properties, including arm length, gene content, CHARM scores, CNA-based net relative fitness values, and common fragile site coverage. Only chromosome arms with ≥10 deletions were included. Each cell is annotated (top to bottom) with the Spearman’s correlation coefficient, p-value, and the fraction of cancer types exhibiting a significant correlation (Spearman’s *p* < 0.05). Only cancer types with ≥ 10 deletions across at least five chromosome arms were analyzed. (E) Pan-cancer distributions of two-hit frequencies across chromosome arms. Distributions for unclassified genes are shown as grey boxplots (with outliers), overlaid with values for individual TSGs (red) and ONGs (blue). TSGs with significant point mutation–deletion associations (Wald test FDR < 0.05, multivariate logistic regression) are outlined in black. (F) Relationship between median pan-cancer two-hit frequencies of unclassified genes and TSGs across chromosome arms. Spearman’s correlation coefficients and p-values are shown in the inset. (G) Relationship between arm-length deletion prevalence and pan-cancer two-hit frequencies for TSGs. Bubble size reflects the total number of deletions. TSGs located on arm-deletion–enriched chromosomes are shown in dark blue, and the rest are shown in light blue. Spearman’s correlation coefficient and p-value are indicated in the inset. (H-I) Co-occurrence and co-deletion of TSG two-hits on chromosomes 3 (H) and 17 (I). Heatmaps (left) show odds ratios for pairwise co-occurrence of TSG two-hit events; asterisks denote significantly co-occurring gene pairs (Fisher’s exact test, FDR < 0.05). Bubble plots (right) show the fraction of co-occurring two-hits driven by co-deletions. Bubble size is proportional to the number of co-occurrence events (minimum ≥ 5 shown). Genes are ordered by genomic position, with p-arm genes highlighted in red.

The above patterns closely follow established aneuploidy landscapes observed independent of TSG alteration status^35^. Consistent with this, arm-length aneuploidy enrichment in TSG two-hit events was broadly comparable to that observed for unclassified genes across cancer types (Spearman’s ρ > 0.6 and *p* < 0.05 in six cancer types, with values close to the diagonal; **Fig. 3C**). This concordance indicates that the two-hit–specific aneuploidy patterns primarily reflect chromosome-intrinsic properties rather than gene-specific effects. We therefore examined how arm-deletion enrichment relates to structural and functional features of the chromosome arms.

In TSG two-hits, arm-deletion enrichment was negatively correlated with chromosome arm length and gene content, consistent with negative selection against loss of large, gene-rich arms to minimize disruption of global gene expression^33^ (ρ = -0.58, *p* = 0.001 and ρ = -0.59, *p* = 0.0009, respectively, Spearman’s correlation; **Fig. 3D**, **Fig. S5C**). We also observed weaker negative correlations with oncogene- and essential gene-specific Charm scores, which capture the net fitness cost of deleting arms enriched for these genes, implying selective constraints imposed by loss of neighboring loci^35^ (ρ = -0.47, *p* = 0.01, and ρ = -0.32, *p* = 0.1, respectively, Spearman’s correlation). Conversely, arm-deletion enrichment was positively correlated with Charm scores combined across TSGs, oncogenes, and essential genes, as well as with net relative fitness estimates derived from copy-number profiles, suggesting a net balance between positive and negative selection^35,36^ (ρ = 0.45, *p* = 0.02, and ρ = 0.41, *p* = 0.03, Spearman’s correlation). Finally, arm-deletion enrichment showed a weak negative correlation with fragile site coverage, suggesting that arms with greater local genomic instability tend to undergo focal rather than arm-length deletions, likely due to increased interstitial breakpoints (Spearman’s ρ = -0.3, *p* = 0.1)^37^. Similar correlation trends were observed in the unclassified control gene set, reinforcing that these patterns are not specific to TSG two-hit events (**Fig. 3D**, **Fig. S5C**).

Taken together, our results suggest that deletion lengths encompassing TSGs in two-hit configurations reflect global aneuploidy patterns shaped by the combined influence of chromosome structure and cumulative selection pressures from neighboring loci. Next, we asked how these aneuploidy patterns relate to the frequency of TSG two-hit events.

### Chromosomal location influences TSG two-hit frequencies

To examine the relationship between deletion patterns and two-hit frequencies, we mapped all genes to their corresponding chromosome arms. We used the unclassified gene set to define background distributions of two-hit frequencies for each arm, assuming these genes evolve under neutral selection. These background distributions varied substantially across the genome, with similar patterns being recapitulated by other negative-control genes (**Fig. 3E**, **Fig. S6A-C**). For example, essential genes showed lower two-hit frequencies than the background, consistent with the lethality of complete gene inactivation, yet their arm-to-arm variation closely mirrored that of the background distributions (Spearman’s ρ = 0.94 and *p* = 1.7e^-18^; **Fig. S6A-B**). Likewise, oncogene two-hit frequencies largely fell within the bounds of the background, showing strong positive correlation with it (Spearman’s ρ = 0.74 and *p* = 6.5e^-7^; **Fig. 3E**, **Fig. S6C**).

While TSGs exhibited higher two-hit frequencies than the background, a similarly strong positive correlation was observed between the two, with chromosome arms harboring TSGs with higher two-hit frequencies also showing elevated background distributions (Spearman’s ρ = 0.8 and *p* = 1.7e^-9^; **Fig. 3E-F**). Further, we observed that TSGs with high two-hit frequencies (≥ 0.4, Wald test FDR < 0.05) were significantly enriched on chromosomes predominantly affected by arm aneuploidies (20 of 27 TSGs; *p* = 0.008, permutation test; see Methods; **Fig. 3E**). This was also reflected in a positive correlation between TSG two-hit frequencies and the prevalence of arm-length deletions in two-hit events affecting each gene (Spearman’s ρ = 0.43 and *p* = 0.0003; **Fig. 3G**).

The concordant variability in two-hit frequencies across gene categories may arise from linked deletion events, highlighting a role for genetic drift in shaping two-hit frequencies. The enrichment of aneuploidies among TSGs with high two-hit frequencies further supports this interpretation, as recurrent TSG two-hit events on these arms would increase the chance of simultaneous alteration of neighboring genes through large, arm-length deletions. Such drift-driven processes could also cause concordance within TSGs, whereby two-hit events in one TSG inflate the two-hit frequency of another. Alternatively, there could be positive selection for the joint inactivation of multiple functionally synergistic TSGs clustered along the same chromosome arm, which would in turn favour aneuploidies over focal deletions. Such cooperative clusters of TSGs could comprise both canonical two-hit genes, as well as haploinsufficient TSGs. Since haploinsufficient TSG clusters are difficult to identify due to the large number of loci affected by hemizygous arm losses, we focused our analysis on identifying clusters of canonical two-hit TSGs.

To this end, we first examined our set of 132 TSGs to identify gene pairs exhibiting significant co-occurrence of two-hit events within the same patients. Among the >8,000 possible combinations, 52 gene pairs showed significant co-occurrence after multiple testing correction (Fishers’ Exact test, FDR < 0.05; see Methods; **Fig. S6D**). Notably, 15 of these pairs involved TSGs located on the same chromosome, representing a significant enrichment over what is expected by chance (*p* < 0.001, permutation test; see Methods). These intra-chromosomal gene pairs were predominantly concentrated on chromosomes 3 and 17. Further inspection revealed that co-occurring two-hits in these pairs were primarily driven by single large deletions spanning both genes (**Fig. 3H-I**, **Fig. S6E-G**).

For example, 82-95% co-occurrence events involving chromosome 3 genes VHL, PBRM1, BAP1 and SETD2 arose through co-deletion (**Fig. 3H**, **Fig. S6F**). These were primarily observed in kidney renal clear cell carcinoma, and involved co-deletion events spanning two to four TSGs. On chromosome 17, co-deletions accounted for 31–69% of co-occurrence events across multiple tumor types, involving genes TP53, CDK12, NF1, and BRCA1 (**Fig. 3I**, **Fig. S6G**). Notably, this also included gene pairs located on opposite arms of the chromosome that were co-deleted through centromere-spanning deletions, such as TP53-NF1 and TP53-BRCA1. Among TP53-containing pairs, co-deletion rates declined with increasing intergenic distance, consistent with the established inverse relationship between CNA length and prevalence (**Fig. 3I**)^33^. In line with our hypothesis, several of these TSG pairs on chromosomes 3 and 17 have been reported to act synergistically in tumorigenesis^38–43^.

Together, these findings indicate that two-hit frequencies are not determined by gene-intrinsic properties alone but are also strongly influenced by chromosomal location and aneuploidy patterns. This is reflected in the mirrored arm-to-arm variability of two-hit frequencies across gene categories, as well as in the frequent co-deletion of multiple TSGs on the same chromosome arm, which results in linked and co-occurring two-hit events.

### TSG all-hit frequencies are preserved despite genome doubling

Aneuploid deletions, found to be the predominant deletion type in TSG two-hits, are frequently elevated in tumors that have undergone whole-genome doubling (WGD), a large-scale genomic alteration in which the entire chromosome set is duplicated one or more times^44–47^. In the TCGA cohort, more than one-third of the samples show evidence of WGD^46^. On one hand, higher deletion rates could increase the probability of acquiring two-hits. On the other hand, WGD raises gene copy number, producing more than two alleles per locus and thereby making complete inactivation more difficult. So far, our analysis of one- and two-hit configurations excluded complex allelic configurations arising from local or global amplifications (see Methods). However, the widespread occurrence of WGD in cancer genomes warrants a more nuanced evaluation.

To assess the impact of WGD on TSG alterations, we redefined two-hit events as “all-hit” events by comparing the total allele count from allele-specific copy number data with the number of mutated alleles estimated from variant allele frequency (VAF) values (see Methods). For each gene, we computed the “all-hit” frequency as the probability of observing an all-hit, conditional on the presence of at least one point mutation, i.e. P (All-hit | Point Mutation) (**Fig. S7A**). All-hit frequencies showed a strong correlation with previously derived two-hit frequencies (Spearman’s ρ = 0.88, *p* < 0.001), underscoring the robustness of this extended framework (**Fig. S7B**).

We next stratified tumors into WGD positive (WGD+) and WGD negative (WGD-) groups and found that pan-cancer TSG all-hit frequencies were highly comparable between the two (Spearman’s ρ = 0.7 and *p* = 3e^-21^; **Fig. 4A**). This concordance persisted at the cancer type level as well, with only rare exceptions (**Fig. S7C**). The same was however not observed for unclassified genes, even when restricting the analysis to those with high (≥ 0.4) two-hit or all-hit frequencies (Spearman’s ρ = -0.42 and *p* = 4.4e^-8^; **Fig. 4B**, **Fig. S7D**). Moreover, both WGD+ and WGD- tumors retained the previously observed hierarchy across mutation classes, with truncating mutations exhibiting higher all-hit frequencies than missense mutations, suggesting that similar selection pressures operate in both genomic contexts (**Fig. S7E**).

**Fig. 4:**
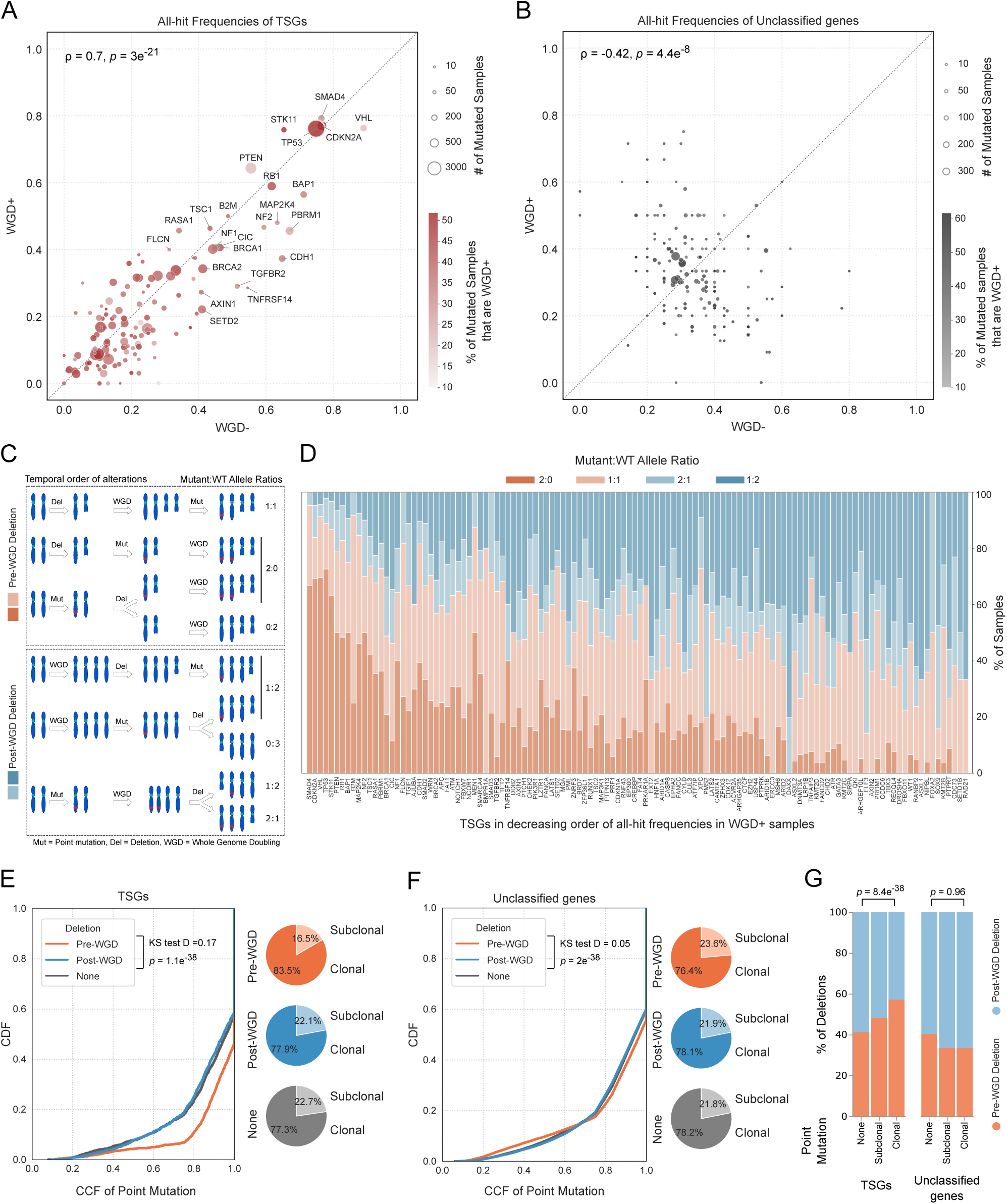
All-hit frequencies and temporal dynamics of alterations in whole genome doubled tumors. (A-B) Pan-cancer all-hit frequencies of TSGs (A) and unclassified genes (B) in WGD+ vs WGD-samples. Only unclassified genes with two-hit or all-hit frequencies ≥ 0.4 across all samples (irrespective of WGD status) are shown. Bubble size denotes the number of samples harboring point mutations, and bubble color indicates the percentage of mutated samples that are WGD+. Insets report Spearman’s correlation coefficients and corresponding *p*-values. (C) Schematic illustrating all possible relative temporal sequences of point mutation (Mut), deletion (Del), and whole-genome doubling (WGD) during clonal evolution, along with resulting end-state mutant:WT allele ratios. (D) Relative prevalence of different mutant:WT allele ratios in TSGs, ordered by decreasing all-hit frequency in WGD+ samples. (E-F) Distributions of cancer cell fraction (CCF) values for point mutations co-occurring with pre-WGD deletions, post-WGD deletions, and no deletions of the remaining allele in TSGs (E) and unclassified genes (F). Insets show Kolmogorov–Smirnov (KS) test D statistics and *p*-values for pre- versus post-WGD CCF distributions. Pie charts indicate the proportions of clonal and subclonal point mutations co-occurring with each deletion category. (G) Prevalence of deletions classified as pre-WGD or post-WGD, stratified by whether the remaining allele carried a clonal, subclonal, or no point mutation in TSGs and unclassified genes. Fisher’s exact test p-values comparing deletion prevalences in cases with clonal versus no point mutation on the remaining allele are shown above each comparison.

Altogether, these results indicate that TSGs acquire all-hits at comparable rates and with similar patterns regardless of genome doubling, consistent with conserved selective landscapes. We next examined the clonal dynamics of TSG alterations to investigate the mechanisms underlying these observations. However, we noted that tumors that had undergone multiple rounds of genome doubling exhibited a downward shift in the distribution of TSG all-hit frequencies, possibly reflecting the increasing difficulty of altering an even larger number of alleles (**Fig. S7F**). Therefore, subsequent analyses were restricted to WGD+ tumors with a single round of genome doubling.

### TSG alterations arise early during clonal evolution

While the persistence of high all-hit frequencies despite WGD could, in principle, arise from multiple independent hits, the parsimonious explanation is that such hits occurred prior to WGD during clonal evolution. In this scenario, mutated alleles are duplicated along with the genome, preserving the effective allelic configuration. To formalise this intuition, we enumerated all possible end-state mutant-to–wild-type (Mutant:WT) allele ratios arising from different relative temporal sequences of point mutation, deletion, and WGD, assuming each event occurs once. Deletions preceding WGD (pre-WGD) yield mutant:WT ratios of 1:1 or 2:0, whereas deletions occurring after WGD (post-WGD) produce ratios of 1:2 or 2:1 (**Fig. 4C**). Ratios such as 0:2 or 0:3 were excluded, as they can be explained without invoking point mutations. When moving from TSGs with high to low all-hit frequencies, we observed a progressive decline in ratios consistent with pre-WGD deletions and a corresponding increase in those indicative of post-WGD deletions, exemplifying the hypothesis that deletions tend to precede WGD in TSGs with high all-hit frequencies (**Fig. 4D**). This trend was driven specifically by a reduction in the 2:0 ratio and a concomitant increase in the 1:2 ratio, suggesting that WGD typically occurs after both point mutation and deletion in TSGs with high all-hit frequencies, and conversely tends to precede both these alterations in those with lower frequencies (**Fig. 4C-D**).

To test this model directly, we classified deletions as pre- or post-WGD depending on whether the minor allele count was reduced to 0 or 1, respectively (see Methods). Point mutations were independently classified as clonal (early) or subclonal (late) based on their cancer cell fraction (CCF) or the proportion of cancer cells harboring the mutation, assuming that earlier events are present in a larger proportion of tumor cells (see Methods). In TSGs, point mutations co-occurring with pre-WGD deletions exhibited significantly higher CCFs, and were therefore more clonal, than those accompanying post-WGD or no deletions (KS D = 0.17; **Fig. 4E**). This pattern was not observed in unclassified genes, which showed similar CCF distributions irrespective of deletion timing (KS D = 0.05; **Fig. 4F**). Conversely, pre-WGD deletions were significantly more prevalent in TSGs when co-occurring with point mutations, particularly with clonal ones (Fisher’s exact *p* = 8.4e^-38^ vs 0.96 for TSGs vs unclassified genes, respectively; **Fig. 4G**). As before, this effect was driven by TSGs with high all-hit frequencies in WGD+ samples (**Fig. S8A**). Collectively, these pan-cancer trends were recapitulated within individual cancer types as well (**Fig. S8B-D**).

Analysis of deletion lengths further revealed that pre-WGD deletions in TSG all-hits were significantly enriched for arm-length events, in line with our earlier observations, whereas post-WGD deletions showed no such enrichment (**Fig. S8E**). Additionally, because deletion timing could only be inferred in WGD+ tumors, we used cancer stage as a proxy for evolutionary time across all samples, assuming earlier stages correspond to earlier time points in tumor evolution. Consistent with early acquisition of alterations, TSGs exhibited similar all-hit frequencies in early versus late cancer stages (Spearman’s ρ = 0.75 and p = 1e^-23^ ; **Fig. S8F**), a pattern not observed for unclassified genes with high (≥ 0.4) two-hit or all-hit frequencies (Spearman’s ρ = -0.3 and *p* = 0.001; **Fig. S8G**).

Overall, these results indicate that TSG point mutations and deletions tend to co-occur early during tumorigenesis, thereby allowing high all-hit frequencies to be preserved despite genome doubling.

## Discussion

Knudson’s two-hit hypothesis has guided our understanding of tumor suppressor gene biology for decades^6^. Although the framework has been substantially refined since its original formulation—incorporating diverse mechanisms of gene inactivation, context-dependent dosage effects, and dominant-negative or gain-of-function modes of action—genome-wide evaluations of how closely individual TSGs adhere to or deviate from this model remain limited^8–10,48–52^. In this study, we systematically charted the two-hit or loss of heterozygosity (LOH) landscape of 132 TSGs through a comprehensive, allele-specific analysis of the TCGA cohort. By defining two hits as point mutation–deletion co-occurrence events and benchmarking their frequencies in TSGs against negative control gene sets, we provide a quantitative, large-scale genomic reinterpretation of a foundational principle in molecular oncology.

We found that two-hit frequency depends strongly on the functional impact of the associated point mutation. Truncating mutations exhibited higher two-hit frequencies than missense mutations, and, for both classes, two-hit frequency scaled with the inferred selection strength of the variants. This relationship, together with the spatial distribution of point mutations along the polypeptide chain, provides the basis for a zygosity-informed approach to variant effect prediction with implications for cancer diagnosis and treatment. TSGs with known dominant mutations exhibited one-hit–specific hotspots, as validated for *TP53* R273 and *PTEN* R130, which accordingly showed comparable transcriptomic impacts between one-hit and two-hit configurations. In contrast, recessive mutations manifested as two-hit–specific hotspots. This was illustrated by the less-characterized missense hotspot mutations in the transcription factor *CIC,* whose transcriptomic profile resembled that of dispersed truncating mutations within the same gene, supporting a loss-of-function effect consistent with prior evidence of impaired DNA-binding activity^30^. The potential of such zygosity-informed strategies for variant interpretation has also been demonstrated by the recent characterisation of variants of unknown significance in KEAP1^10^. Future studies that integrate mutation positions from three-dimensional protein structures and interaction interfaces are likely to further strengthen such approaches.

Our analysis identified chromosomal location as another important determinant of two-hit frequency, with TSG-specific frequencies closely mirroring background trends derived from unclassified genes located on the same chromosome arm. This pattern likely reflects chromosome-intrinsic aneuploidy biases, whereby the probability of deletion, and consequently of a two-hit event, depends on the structure and genetic composition of the chromosome arm. For instance, reduced two-hit frequencies in some TSGs may arise from negative selection against deletions that simultaneously disrupt neighboring essential genes or oncogenes. Conversely, deletions overlapping physically linked functionally synergistic TSGs may confer a selective advantage by enabling their co-inactivation through fewer mutational events. We identified several putative synergistic TSG pairs, predominantly on chromosomes 3 and 17, in which two-hit events frequently co-occurred in the same samples via shared deletions. Notably, both chromosomes exhibited arm-length deletions despite moderate essentiality scores, suggesting that the fitness gains from co-inactivation of synergistic TSGs can outweigh opposing negative selection pressures from adjacent loci (**Fig. 3A**, **Fig. 3H-I**, **Fig. S5C**). Nevertheless, our analysis likely underestimates the prevalence of such co-deleted TSG pairs. By focusing on significantly co-occurring two-hit events, we excluded TSGs inactivated through single (hemizygous) or double (homozygous) deletions, and our search was further restricted to a highly conservative set of high-confidence TSGs. For example, *WRN* was the only TSG from chromosome 8p included in our analysis^36^. However, evidence from murine models and patient data suggests that 8p loss disrupts multiple candidate TSGs that cooperatively promote hepatocellular carcinoma progression^53^.

Additionally, we investigated the implications of polyploidy arising from WGD for TSG inactivation. We found that the frequencies of TSG “all-hit” events—two-hit equivalents in polyploid contexts with more than two alleles per gene—are largely similar between WGD+ and WGD- cancers. Thus, although certain TSGs such as *TP53* are mutated more frequently in WGD+ tumors, the probability of losing the remaining alleles is comparable to that in WGD- tumors^46,54^. Analysis of end-state allelic configurations and mutation clonal dynamics suggests that point mutations and deletions frequently co-occur prior to WGD during clonal evolution, allowing these alterations to be duplicated and preserved through genome doubling. While WGD is already known to induce genomic instability and elevate global aneuploidy levels, these changes typically occur after the doubling event^44,46,47^. However, for TSGs, deletions preceding WGD provide the parsimonious route to effective LOH across multiple alleles. This interpretation is supported by prior evidence of positive selection for pre-WGD LOH of TSGs in lung cancer, as well as by recent reports showing preferential pre-WGD deletions on several chromosome arms harboring key TSGs (including 17p, 9p, and 8p), in contrast to predominantly post-WGD loss elsewhere^55,56^. Consistent with this model, additional studies indicate that WGD typically occurs as an intermediate-to-late event during cancer evolution across multiple cancer types^57,58^. However, this model does not appear to extend to higher ploidy states arising from multiple rounds of WGD, where we observed a deviation in the distribution of TSG all-hit frequencies (**Fig. S7F**), warranting further investigation into the underlying mutational dynamics and their implications. Additionally, we acknowledge that post-WGD deletions in TSGs may retain functional relevance through dosage-sensitive mechanisms, even when not resulting in all-hit configurations^59,60^.

The persistence of high all-hit frequencies despite WGD may reflect strong selection to maintain TSG LOH even in later stages of tumor evolution. Alternatively, TSG LOH may be most critical during early tumorigenesis, with WGD arising later as a consequence of tumor progression or genomic instability induced by early LOH events. In this case, stable all-hit frequencies would represent a byproduct of shifting selective pressures rather than ongoing selection. Consistent with this interpretation, TSG all-hit frequencies remain stable across cancer stages irrespective of WGD status. Moreover, prior studies indicate that many driver events can arise years before clinical diagnosis and are often under selection even in healthy tissues^57,61–63^. If this also applies to TSG LOH, assessing TSG allelic status could provide greater clinical utility for cancer screening and monitoring than hotspot mutation analysis alone.

This study focused exclusively on two-hit or all-hit events arising from coding-region genetic alterations. Other modes of gene inactivation, including regulatory-region alterations and epigenetic silencing, were beyond the scope of this study but merit further investigation. Besides, while point mutation-deletion combinations can reasonably be assumed to affect different alleles, multiple point mutations could not be phased to distinct haplotypes and were therefore treated as one-hit events, potentially leading to modest underestimation of the computed two-hit frequencies. Finally, we did not account for the influence of pre-existing genetic alterations in other genes within the same cell on the probability of acquiring TSG alterations. Prior work has shown that epistatic interactions within interaction networks can modulate biallelic alteration rates of TSGs, revealing additional layers of complexity⁵.

In summary, our analysis provides a quantitative framework for interpreting tumor suppressor gene alterations across molecular and temporal dimensions in cancers. By systematically characterizing the genomic events underlying loss of heterozygosity, we delineate how the impact of point mutations, chromosomal context, and polyploidy jointly shape the probability of acquiring two hits across TSGs. Together, these insights extend the classical two-hit paradigm and provide a more comprehensive understanding of cancer driver gene behavior during clonal evolution, laying the foundation for improved cancer diagnostics and precision medicine.

## Methods

### Allelic configurations of genes

Allelic configurations of genes were determined in 8,958 patients across 33 cancer types in The Cancer Genome Atlas (TCGA) by integrating data for copy number profiles with point mutations (SNVs, short INDELs, small sequence alterations). Somatic point mutation data were obtained from Ellrott et al. (2018), as compiled by the MC3 Working Group^12^. Low-quality and hypermutator samples were excluded following the criteria outlined by Bailey et al. (2018)^64^. Somatic variants were supplemented with non-synonymous pathogenic germline variants from TCGA, as reported by Huang et al. (2018)^65^.

Allele-specific copy numbers were derived from Affymetrix SNP array profiles of TCGA samples using the ASCAT pipeline^13,66^. ASCAT genomic intervals were mapped to the curated point mutations in each patient to infer local copy number status based on major and minor allele counts. Loci with a minor allele count of zero were considered to have copy number loss or deletion. Point mutations with a total copy number of zero (major = minor = 0) were excluded as likely technical artefacts. This resulted in 1,228,437 copy-number–annotated variants, including missense (778,124), nonsense (57,206), frameshift indels (45,202), splice-site (21,602), in-frame indels (6,635), translation start-site (1,370), and silent (318,298) mutations. The vast majority were somatic variants (1,227,251), with the remainder comprising pathogenic (369), likely pathogenic (356), and prioritised variants of uncertain significance (461) of germline origin. Annotated variants were used to assign allelic configurations to genes, including those harboring one or multiple point mutations (minor > 0), and those with both point mutation and deletion (minor = 0).

For genes without variants, local copy number status was assigned by mapping gene coordinates to ASCAT genomic intervals overlapping with at least two-thirds of the gene length. This enabled classification of genes with deletion of one (major > 0, minor = 0) or both (major = minor = 0) alleles, or with neither deletion nor point mutation (minor > 0). Gene coordinates were obtained from Ensembl BioMart^67^.

### Driver and non-driver gene sets

Multiple databases were used to curate lists of driver (TSG, ONG) and non-driver (essential, non-essential, unclassified) genes used in our analyses. Driver genes were compiled from Bailey et al. (2018) (included if classified as a driver in at least one cancer type without conflicting TSG vs ONG annotations in others), IntOgen (2020) catalog of driver genes and Cosmic Cancer Gene Census (2024)^4,64,68,69^. A high-confidence set of TSGs was derived from genes annotated as a TSG in at least two of the three databases, with no conflicting classification in the third; high-confidence ONGs were identified using the same criterion. Essential genes were defined as those annotated as essential in at least two of the following sources: Caitlin et al. (2020), CRISPR KO common essential genes, and Achilles essential positive control genes from the DepMap 24Q2 Public Release (https://plus.figshare.com/articles/dataset/DepMap_24Q2_Public/25880521)^70–73^. Non-essential genes were derived from the Achilles essential negative control genes from the DepMap project^72,73^. Genes not classified in any of the source gene lists were labelled as unclassified genes. Across all classifications, only protein-coding genes having somatic non-synonymous mutations in at least five TCGA patients were considered. Genes located on sex chromosomes were excluded from the analysis as they are haploid in males and undergo X-inactivation in females. This resulted in 132 TSGs, 115 ONGs, 1312 essential genes, 703 non-essential genes and 12536 unclassified genes. Among the protein-coding genes present in the source databases of driver and essential genes, the 2495 that were not included in our curated gene sets were labelled ambiguous.

### One-hit and two-hit classification

To quantify the co-occurrence of point mutations and deletions, each driver or non-driver gene carrying a point mutation was annotated as either “one-hit” or “two-hit” in each patient. A gene was considered “one-hit” if it harbored at least one coding non-synonymous point mutation in the absence of copy number loss or deletion (minor > 0). This configuration corresponds to a point mutation on one allele with the other allele remaining wild type, but also includes a minority of cases with multiple point mutations whose haplotypes could not be phased. Conversely, genes with a coding non-synonymous mutation that also exhibited copy number deletion (minor = 0) were classified as “two-hit”, indicating a point mutation on one allele and deletion of the other. Genes without any point mutations were similarly classified as having a deletion (minor = 0) or not (minor > 0).

This initial annotation was limited by the assumption that there is no local or global chromosomal amplification affecting the gene locus in tumor cells. To refine this classification, the number of mutated chromosomal copies, also referred to as the multiplicity of the mutation or *n_chr_*, was estimated following the method of Letouzé et al. (2017)^74^, computed as the integer closest to:

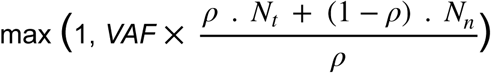

where *VAF* is the variant allele fraction (i.e., the fraction of sequencing reads at the locus supporting the variant), ρ is the tumor purity (obtained from ASCAT), *N_t_* is the total copy number at the locus in tumor cells (sum of major and minor allele counts), and *N_n_* is the copy number in normal cells (considered diploid, i.e., 2). Among the initially defined two-hit cases (minor = 0), those with *n_chr_* < *N_t_* were excluded, as such low multiplicity contradicts the expected deletion of the second allele. Similarly, among initially defined one-hit cases (minor > 0), those with *n_chr_* ≥ *N_t_* were excluded, as this indicates unexpectedly high mutation multiplicity. Additionally, one-hit cases in which the number of wild-type allele copies (*N_t_* − *n_chr_*) exceeded the number of mutant copies (*n_chr_*) were excluded due to the low relative dosage of the mutant allele. For genes without any point mutations, those with and without a deletion were only considered for further analysis if their total copy number (*N_t_)* was 1 and 2, respectively.

#### Statistical evaluation of two-hit frequencies

The two-hit frequency of each gene was defined as the proportion of patients with point mutations who also exhibited a two-hit event:

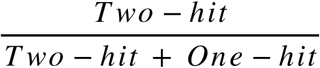

This is equivalent to the conditional probability of a deletion given the presence of a point mutation, i.e., *P (Deletion | Point Mutation)*. Pan-cancer or cancer-type–specific two-hit frequencies were calculated only for genes mutated in at least five patients across or within cancer types, respectively. For TSGs, this resulted in 132 pan-cancer and 790 cancer-type–specific two-hit frequency values.

To assess the significance of these frequencies, we modeled deletion as a function of point mutation status using multivariate logistic regression, with tumor mutation burden (TMB) and fraction of genome altered (FGA) as additional covariates to account for genome-wide point mutation and copy number burden:

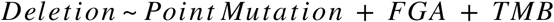

TMB and FGA data were downloaded from https://gdc.cancer.gov/about-data/publications/panimmun^75^. The model was fit independently for each TSG at both the pan-cancer and cancer type levels. To mitigate bias from small sample sizes or skewed data distributions, particularly within cancer type stratifications, we employed Firth’s penalized likelihood correction as previously done in Zucker et al. (2025)^10^. Statistical significance was evaluated from the Firth logistic regression coefficient estimates using two-sided Wald tests. This yielded interpretable *p*-values for the mutation coefficient across 780 of the 790 TSG-cancer-type pairs tested. For VHL in KIRC, the mutation predictor resulted in a very large coefficient with a narrow confidence interval that excluded zero, indicating a strong association with deletion. However, the *p*-value could not be reliably estimated, possibly due to quasi-complete separation, even with Firth’s correction. Obtained *p*-values were adjusted for multiple hypothesis testing using the Benjamini–Hochberg false discovery rate (FDR) correction. For cancer-type–specific analyses, this correction was applied separately within each cancer type. TSGs were considered significant if their FDR-adjusted *p*-values were less than 0.05.

### Estimation of selection coefficients

Selection coefficients of point mutations were estimated at the pan-cancer level using the *dNdScv* R package^13^. Samples harboring more than 1000 coding mutations were excluded, and analyses were restricted to a maximum of three mutations per gene per sample. Maximum-likelihood estimates of dN/dS ratios were obtained along with 95% confidence intervals, likelihood ratio test *p*-values, and Benjamini-Hochberg–adjusted FDR values for truncating and missense mutations in one-hit and two-hit configurations. This was done for individual genes, as well as globally across all TSGs, TSGs stratified by high versus low pan-cancer two-hit frequency, and across additional driver and non-driver gene sets.

### Functional classification of point mutations

Variant classification annotations provided by TCGA were used to compute mutation-class–specific two-hit frequencies. Mutations predicted to cause stop-gain, splice site, frameshift, or start-loss effects were grouped as truncating. Variants annotated as missense and synonymous were categorised accordingly, while in-frame indels were excluded from classification due to their uncertain functional impact. When multiple variants occurred within the same gene in a given patient, truncating mutations were assigned highest priority; missense mutations were considered only in the absence of truncating or in-frame variants; and synonymous mutations were assigned only when no non-synonymous variants were present. Missense mutations were further stratified by predicted functional impact using scaled (PHRED-like) CADD (Combined Annotation Dependent Depletion) scores obtained from version 1.6 of the CADD database (https://cadd.gs.washington.edu/download)^14^. When multiple missense variants were present in a gene, the variant with the highest CADD score was prioritised. Variants with CADD scores above the third quartile were classified as high impact, whereas those below the first quartile were classified as low impact. Two-hit frequencies were then calculated as described above, but separately for each mutation category, yielding conditional probabilities of observing a deletion given the presence of point mutation from a specific functional category.

### Spatial distribution of point mutations

Mutation positions were compared between one-hit and two-hit configurations using two complementary statistical approaches: the Kolmogorov–Smirnov (KS) test, which assesses differences between cumulative distributions of mutation positions, and the Earth Mover’s Distance (EMD) test, which quantifies global shifts in mutation density along the protein sequence. For the KS test, mutation positions were treated as one-dimensional distributions, and the KS statistic was computed between one-hit and two-hit groups. Empirical *p*-values were obtained by comparing the observed statistic to permutation-based null distributions generated by repeatedly shuffling mutation labels (10,000 iterations) between the two groups.

For the EMD analysis, mutation positions were represented as weighted histograms spanning the length of the protein containing the point mutations, and the first-order EMD or Wasserstein distance was computed between the two configurations. To account for differences in protein length and mutation burden, the observed EMD was normalized by the theoretical maximum possible distance, defined as the distance obtained if all mutations in one group occurred at the N-terminus and all in the other group occurred at the C-terminus. Null distributions of normalized EMD values were generated by permutation (10,000 iterations) while preserving group sizes, and empirical *p*-values were calculated as the proportion of permuted values greater than or equal to the observed statistic.

Both tests were performed separately for truncating and missense mutations. To minimise biases arising from sparse data, analyses were restricted to TSGs with at least five mutations in both one-hit and two-hit configurations across all cancer types. Further, to increase robustness, we focused on genes with recurrent mutation positions or hotspots in either configuration, defined as sites mutated in at least five patients. This yielded 39 TSGs with pan-cancer hotspot mutations, of which 25 also harbored hotspots within at least one cancer type. Results corresponding to the mutation class of the most frequent hotspot were reported for each gene. Genes with a *p*-value < 0.1 in either test were considered to exhibit significantly different mutation position distributions between one-hit and two-hit configurations. Statistical results were corroborated with visual representations of mutation positions and frequencies. Protein domain annotations for the lollipop plots were obtained from Pfam and Interpro^76,77^.

### Enrichment of hallmark pathways

Transcriptomic signatures of mutant and wild-type samples were assessed by computing single-sample gene set enrichment scores using the GSVA R package^78^. Enrichment analyses were performed on gene expression count matrices separately for each cancer type, applying the GSVA method with a Poisson kernel to 50 Hallmark gene sets from MSigDB^79^. Pathways dysregulated between samples harboring point mutations and those that were wild-type or carried a single-allele deletion were identified by comparing GSVA enrichment scores using two-sided Mann–Whitney U tests, with *p*-values adjusted for multiple testing using the Benjamini–Hochberg procedure. Differences in pathway enrichment among samples with point mutations (one-hit vs two-hit; two-hit truncating vs two-hit missense) were evaluated using a two-sided Mann–Whitney U test. For CIC, missense mutations between protein positions 200–230 or beyond position 1500 were considered hotspots.

### Determining deletion lengths and aneuploidies

To determine the lengths of deletions associated with two-hit events, we first identified contiguous deletion segments in each of the 8,958 TCGA patients. Adjacent ASCAT intervals with minor allele copy number equal to zero were merged if separated by gaps of no more than 1,000 bp, a threshold chosen based on the elbow of the inter-interval gap size distribution. These merged intervals were treated as deletions of the same haplotype. The genomic coordinates of each merged segment defined the location and length of a unique deletion. For isolated intervals with minor allele copy number equal to zero, the ASCAT interval itself was considered a unique deletion.

Deletion lengths were normalized by the length of the corresponding chromosome arm. For deletions spanning the centromere, normalized length was calculated as the sum of the deleted fractions of the p and q arms, following Beroukhim et. al, (2010). Chromosome arm lengths were derived from cytoband coordinate annotations downloaded from the UCSC Genome Browser^80^. Deletions covering ≥ 70% of a chromosome arm were classified as arm-length (aneuploid), those covering 51–69% as large deletions, and those covering ≤ 50% as focal. Centromere-spanning deletions were classified as chromosomal deletions if they removed ≥ 70% of both arms. For the remaining centromere-spanning events, deletions in which the overlap differed by more than 20% between arms were assigned to the arm with greater overlap and classified as focal, large, or arm-length using the same thresholds. Deletions removing similar proportions of both arms (≤ 20% difference) could not be unambiguously assigned and were left unclassified.

Acrocentric chromosomes (chromosomes 13, 14, 15, 21, and 22) were excluded only in chromosome arm-level analyses. When calculating the percentage of arm-length deletions per chromosome arm, only arms with at least 10 total deletions (focal + arm-length) were included, both in pan-cancer and cancer-type–specific analyses. For unclassified genes, two-hit–associated deletions were included only if the deletion overlapped a point mutation in that gene (after filtering for amplification bias as described above) and did not overlap any non-synonymous coding-region point mutations in TSGs, oncogenes, essential genes, or ambiguous genes (prior to filtering for amplification bias).

### Chromosome arm properties

Lengths of chromosome and chromosome arms were derived from cytoband coordinate annotations downloaded from the UCSC Genome Browser^80^. Charm scores, which quantify the net fitness effect of a chromosome arm based on the density and functional impact of the essential and cancer driver genes it contains, were obtained from Davoli et al. (2013)^35,36^. Net relative fitness estimates derived from copy-number alteration profiles were obtained from Shih et al. (2023)^35,36^. Coordinates of common fragile sites (CFS) were obtained from Kumar et al. (2019)^37^. Fragile site coverage of each chromosome arm was calculated as the fraction of the arm that overlapped annotated CFS regions.

### Co-occurrence and Enrichment Analyses

Pairwise co-occurrence of TSG two-hit events was evaluated using 2×2 contingency tables across all gene pairs, with significance assessed by two-sided Fisher’s exact tests. Odds ratios were computed with a small continuity correction (0.001 added to all cells) to avoid instability due to zero counts, and *p*-values were adjusted using the Benjamini–Hochberg method.

Permutation testing was used to evaluate (i) whether TSG pairs exhibiting significant co-occurrence of two-hit events were located on the same chromosome more frequently than expected by chance, and (ii) whether TSGs with high two-hit frequencies were significantly enriched on chromosomes predominantly affected by arm aneuploidies. For the first, we generated a null distribution by randomly sampling an equal number of gene pairs from all possible TSG pairs and counted the number of intra-chromosomal pairs in each iteration (10,000 permutations). For the second analysis, we repeatedly sampled random TSG sets of identical size and recalculated the chromosomal overlap in each iteration (10,000 permutations). In both analyses, empirical one-sided p-values for enrichment were calculated as the proportion of permutations yielding values greater than or equal to the observed statistic, with a pseudocount of +1 applied to both the numerator and denominator.

### All-hit frequencies

Each driver or non-driver gene carrying a point mutation was annotated as either “all-hit” or “some-hit” in each patient by comparing the number of mutated alleles (multiplicity or *n_chr_*) with the gene’s total copy number (*N_t_*) or the sum of the major and minor allele counts. Genes with *n_chr_ ≥ N_t_* were classified as all-hits, indicating that all non-deleted alleles carried a point mutation. Genes with *n_chr_ < N_t_* were classified as some-hits, indicating the presence of both point-mutated and wild-type (WT) alleles. The all-hit frequency of each gene was defined as the proportion of patients with point mutations who also exhibited an all-hit event:

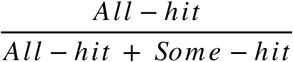

This corresponds to the conditional probability of observing an all-hit event given a point mutation, i.e., *P (All-hit | Point Mutation)*.

Samples with whole genome doubling (WGD) were identified based on the percentage of the autosomal genome with major copy number ≥ 2 as annotated in Bielski et al. (2018)^81^. All-hit frequencies were calculated across all samples, as well as separately for WGD+ and WGD- samples. Samples with zero genome doubling events were classified as WGD-, whereas those with one or more genome doubling events were classified as WGD+. All-hit frequencies were also calculated separately for early-stage (I–II) versus late-stage (III–IV) tumors, where cancer stage information was available (21 cancer types). Pan-cancer and cancer-type–specific all-hit frequencies were calculated only for genes mutated in at least five patients across or within cancer types, respectively.

### Relative timing of deletions

Deletion events were timed relative to WGD in patients with one genome doubling event by assuming the most parsimonious sequence of alterations as described in Zack et al. (2013)^82^. Briefly, copy-number changes were modeled starting from one copy each of the major and minor alleles. ASCAT intervals with major = 2 and minor = 2 were interpreted as having undergone only WGD, with no accompanying deletion. Intervals with major = 2 and minor = 1 were annotated as deletions occurring after WGD (post-WGD), whereas intervals with major = 2 and minor = 0 were annotated as deletions occurring before WGD (pre-WGD). Intervals with major allele copy numbers greater than or less than two were inferred to have undergone additional local amplification or deletion, respectively, and were therefore excluded from the timing analyses due to ambiguity. Lengths of pre-WGD and post-WGD deletions were calculated and normalized as described before: pre-WGD deletion boundaries were defined based on contiguous stretches of ASCAT intervals with major = 2 and minor = 0, while post-WGD deletion boundaries were defined from contiguous stretches with major = 2 and minor = 1. Each unique deletion event was counted only once when comparing the prevalence or lengths of pre-WGD versus post-WGD deletions.

### Clonality of point mutations

The cancer cell fraction (*CCF*), defined as the proportion of cancer cells harboring a given point mutation, was estimated using the approach described in McGranahan et al. (2015)^54^. For each mutation, the expected variant allele frequency (*eVAF*), or the fraction of sequencing reads supporting the variant, was computed over a uniform grid of candidate *CCF* values as a function of tumor purity (ρ), variant multiplicity (*n_chr_*), total copy number in tumor cells (*N_t_* = sum of major and minor allele counts), and the total copy number in normal cells (*N_n_* = 2, assuming diploidy) as follows:

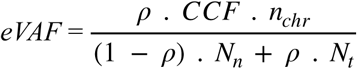

For each candidate *CCF* value, the likelihood of observing *k* alternate reads out of a total sequencing depth *D* was computed under a binomial model:

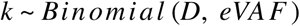

This likelihood function was normalized to yield a posterior distribution over *CCF* values, assuming a uniform prior. The maximum a posteriori estimate was assigned as the *CCF* of the mutation, and a 95% credible interval (CI) was derived from the posterior distribution. Mutations with an upper CI bound exceeding 1 were classified as clonal, whereas the remaining mutations were classified as subclonal. Patients in whom > 95% of mutations were classified as subclonal were excluded, suspecting poor read quality. For genes or deletion segments harboring multiple point mutations within the same patient, the mutation with the highest CCF was prioritised and retained for downstream analyses.

## Acknowledgements

We acknowledge the funding support from the Department of Atomic Energy, Government of India, under Project Identification No. RTI 4006 and intramural funds from NCBS-TIFR. RS acknowledges support from the DBT/Wellcome Trust India Alliance Fellowship [grant number IA/I/20/1/504928]. NM would like to acknowledge funding support from the Shyama Prasad Mukherjee Fellowship (SPMF) awarded by the Council for Scientific and Industrial Research, Ministry of Science & Technology, Government of India. We thank Deepa Agashe, Dimple Notani, Shaon Chakrabarti, Anurag Kumar Singh, and members of RS lab, especially Surya Sarathi Das and Disha Kshirsagar, for their feedback and suggestions on this manuscript.

## Author contributions

NM and RS conceived and designed the study. NM performed the data curation and analysis, interpreted the results and wrote the manuscript with inputs from RS. RS supervised the study. All authors read and approved the final manuscript.

## Supplementary Figures

**Fig. S1:**
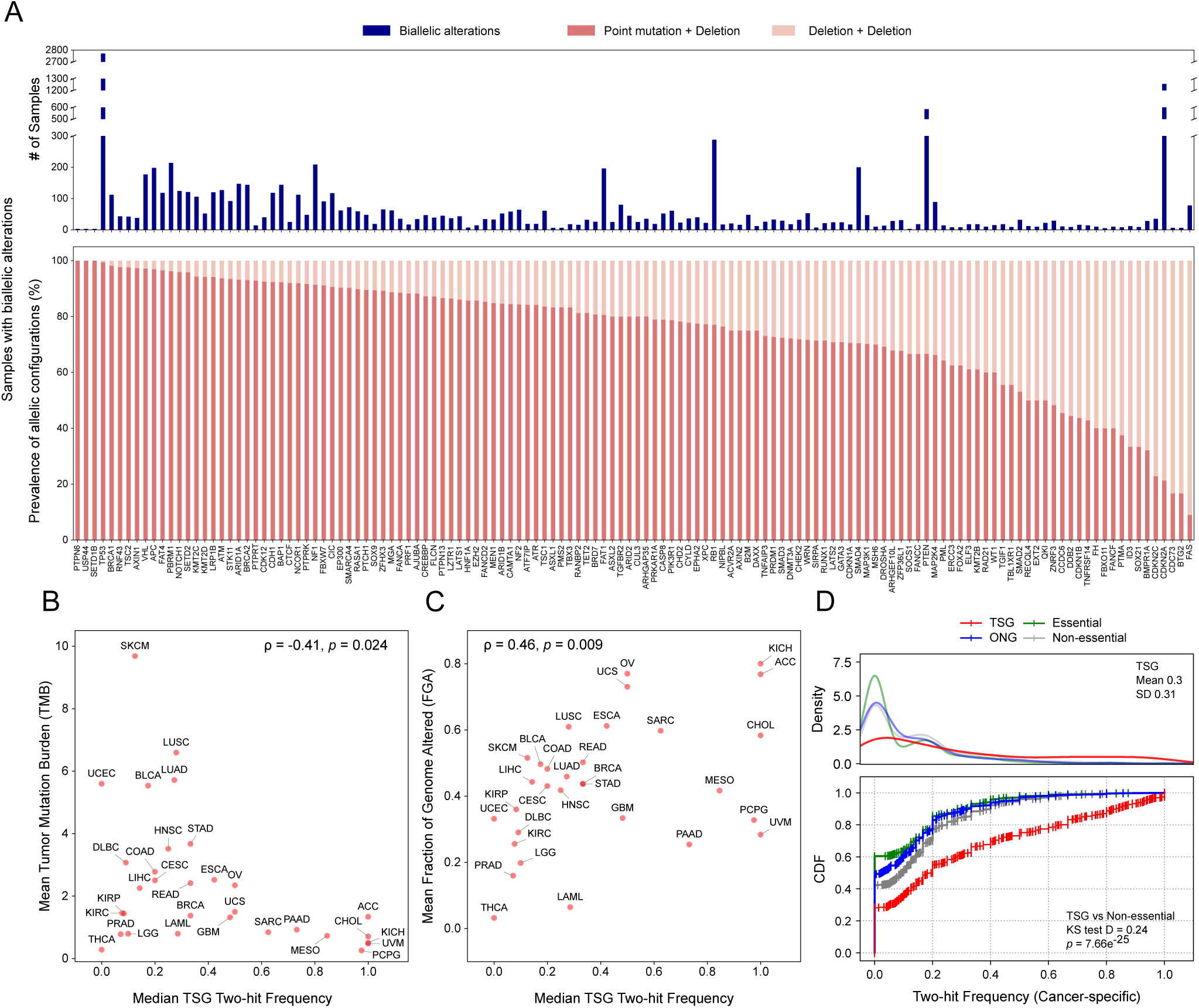
Biallelic alterations in TSGs and two-hit frequencies. (A) Number of samples with biallelic alterations in TSGs (top) and the relative prevalence of underlying allelic configurations: Point Mutation + Deletion and Deletion + Deletion (bottom). (B-C) Relationship between TSG two-hit frequencies and TMB (tumor mutation burden) (B) or FGA (fraction of genome altered) (C) across cancer types. Insets show the Spearman’s correlation coefficients and *p*-values. (D) Kernel Density Estimations or KDEs of cancer-type–specific two-hit frequencies across gene categories, with insets showing the mean and standard deviation of the TSG distribution (top). Cumulative Distribution Functions or CDFs of cancer-type–specific two-hit frequencies across gene categories, with insets displaying the Kolmogorov-Smirnov (KS) distance and p-value comparing TSGs and non-essential genes (bottom).

**Fig. S2:**
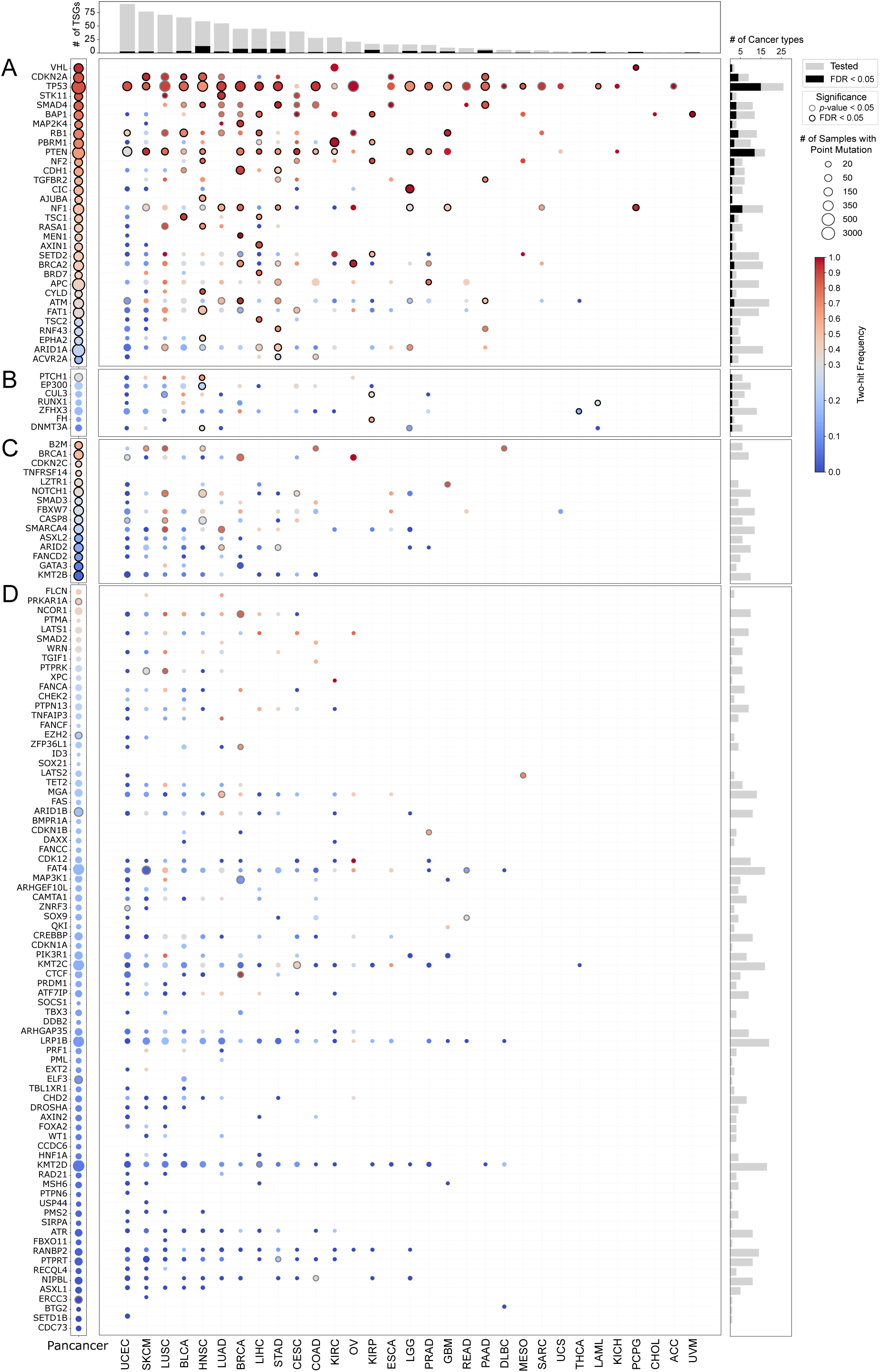
Cancer-type–specific two-hit frequencies of TSGs. Bubble plot showing two-hit frequencies for TSG–cancer-type pairs. Bubble color represents two-hit frequency value, with darker red indicating higher frequencies and darker blue indicating lower frequencies. Bubble size corresponds to the number of samples with point mutations. Only TSG–tumor pairs with ≥ 5 samples harboring point mutations are shown. Pairs with statistically significant point mutation–deletion associations are outlined in black and grey (Wald test FDR < 0.05 and *p* < 0.05, respectively, multivariate logistic regression). TSGs are grouped into the following groups (top to bottom): (A) significant at the pan-cancer level and in at least one cancer type, (B) significant only in specific cancer types, (C) significant only at the pan-cancer level, and (D) not significant in either context. Bar plots on the top show the number of TSGs tested (grey) and significant (black; FDR < 0.05) for each cancer type. Bar plots on the right show the number of cancer types tested (grey) and significant (black; FDR < 0.05) for each TSG.

**Fig. S3:**
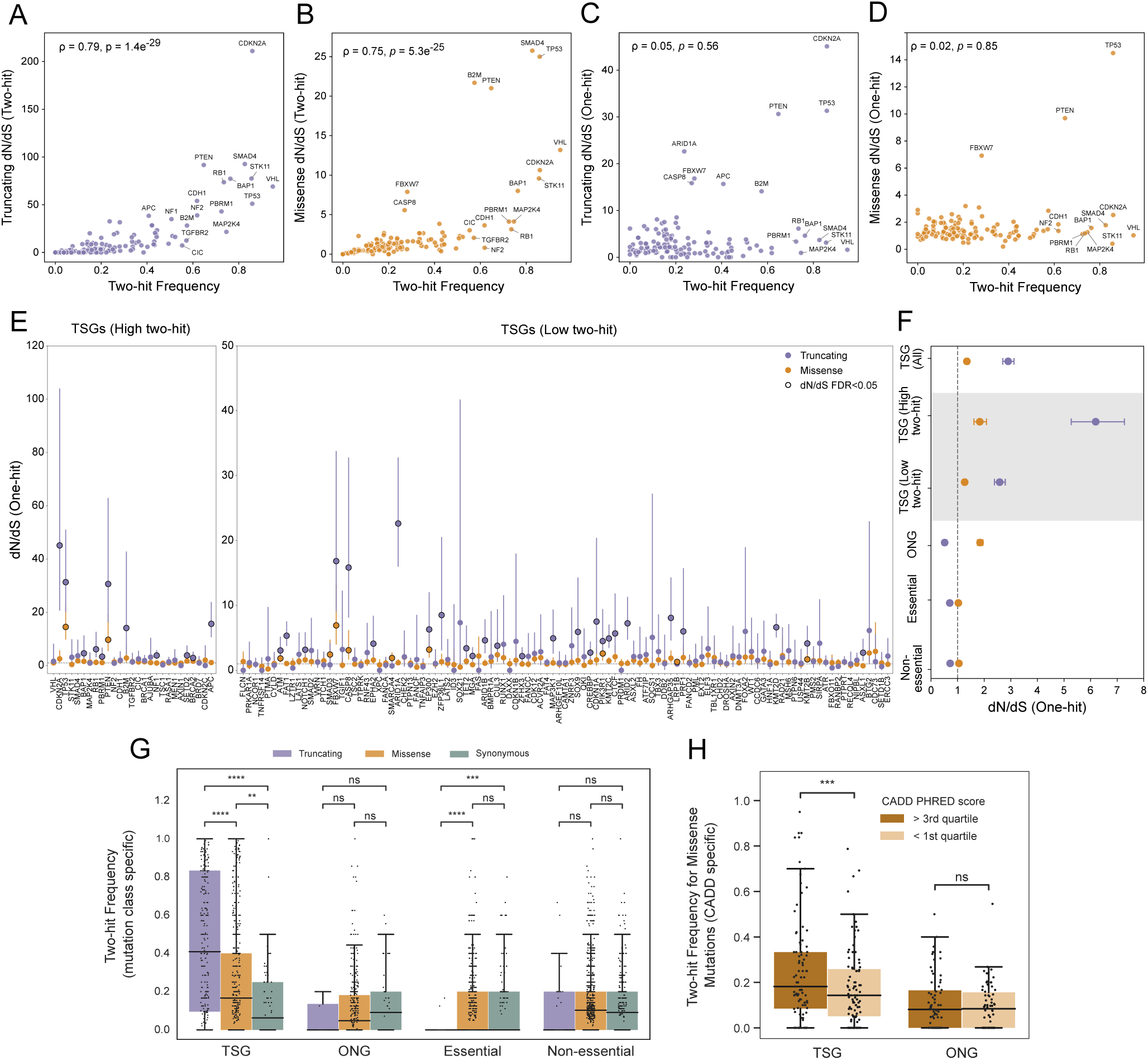
Functional impact of point mutations shapes two-hit frequencies. (A-D) Scatter plots illustrating the relationship between pan-cancer two-hit frequencies of TSGs and dN/dS ratios, including two-hit–specific dN/dS ratios for truncating (A) and missense (B) mutations, and one-hit–specific dN/dS ratios for truncating (C) and missense (D) mutations. Insets report Spearman’s correlation coefficients and corresponding p-values. (E) One-hit–specific dN/dS ratios for truncating (purple) and missense (orange) mutations in TSGs, ordered along the x-axis by decreasing pan-cancer two-hit frequency. TSGs with high two-hit frequencies (≥ 0.4, Wald test FDR < 0.05) are shown separately (left). Horizontal dashed line represents dN/dS value of 1. dN/dS ratios significantly different from 1 (FDR < 0.05, Likelihood ratio test) are outlined in black. (F) Global one-hit–specific dN/dS ratios for truncating (purple) and missense (orange) mutations in TSGs, ONGs, essential genes and non-essenial genes. Global estimates calculated separately for TSGs with high and low two-hit frequencies are also shown. (G) Distributions of cancer-type–specific two-hit frequencies of different gene categories, computed separately for truncating, missense and synonymous. Asterisks denote statistical significance based on paired Wilcoxon signed-rank tests: *p* > 0.05 (ns), ≤ 0.05 (*), ≤ 0.01 (**), ≤ 0.001 (***), ≤ 0.0001 (****) (H) Distributions of pan-cancer two-hit frequencies for missense mutations in TSGs, computed separately for missense mutations with high (> 3rd quartile) and low (< 1st quartile) CADD PHRED scores. Asterisks denote statistical significance based on paired Wilcoxon signed-rank tests: *p* > 0.05 (ns), ≤ 0.001 (***)

**Fig. S4:**
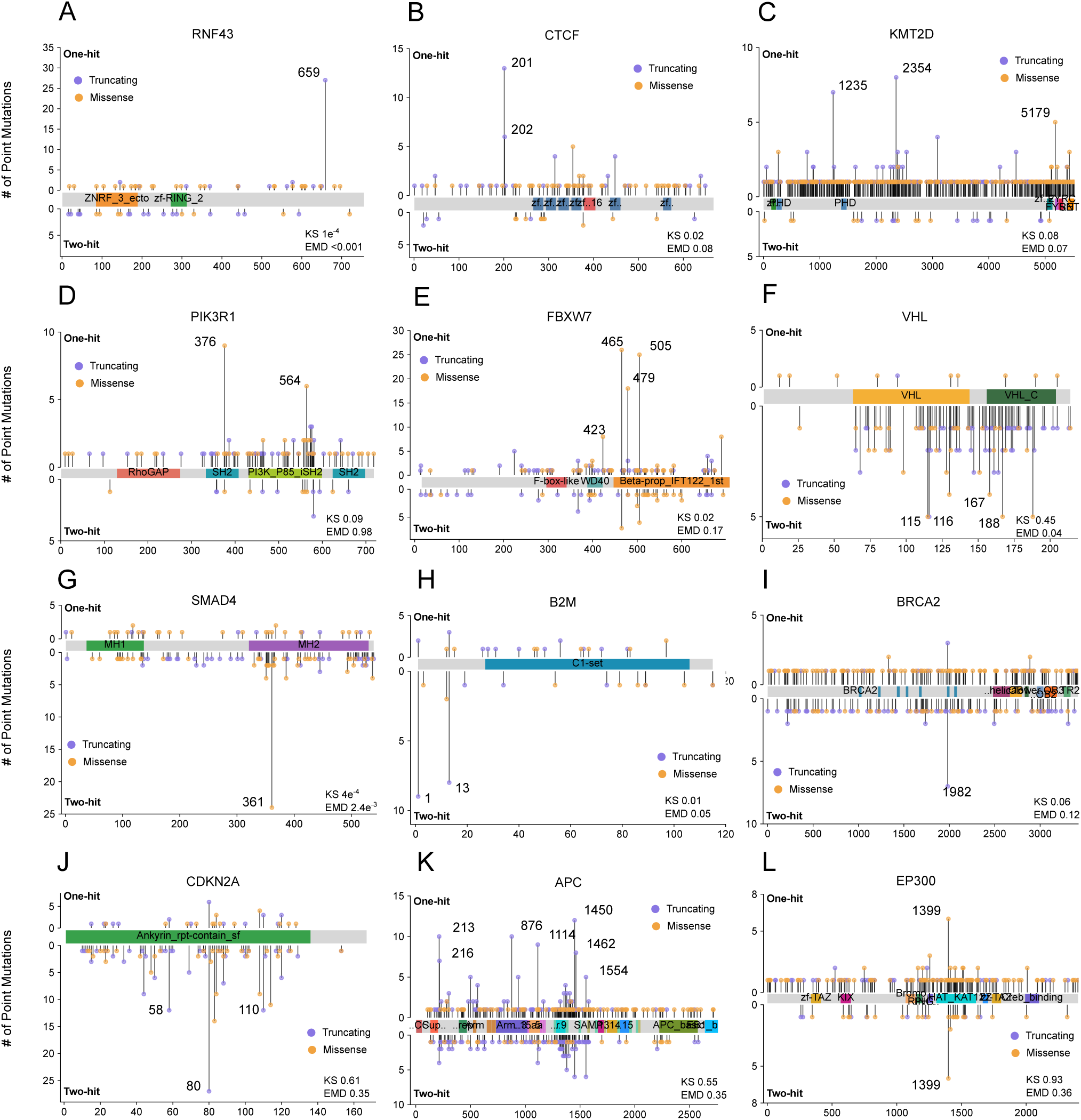
Spatial mutation patterns across one- and two-hit configurations. Lollipop plots showing the distribution of point mutation positions along the linear polypeptide for one-hit and two-hit configurations in representative TSGs. X-axis represents amino acid position. Missense mutations are shown in yellow and truncating mutations in purple. Recurrent single-base hotspots (mutated in ≥ 5 patients) are annotated. Insets show p-values comparing one-hit and two-hit mutation position distributions for the mutation class of the most frequent hotspot in that gene, using Earth Mover’s Distance (EMD) and Kolmogorov–Smirnov (KS) tests. Examples illustrate recurrent point mutations unique to one-hit (A-E), unique to two-hit (F-I), and shared between one- and two-hit (J-L).

**Fig. S5:**
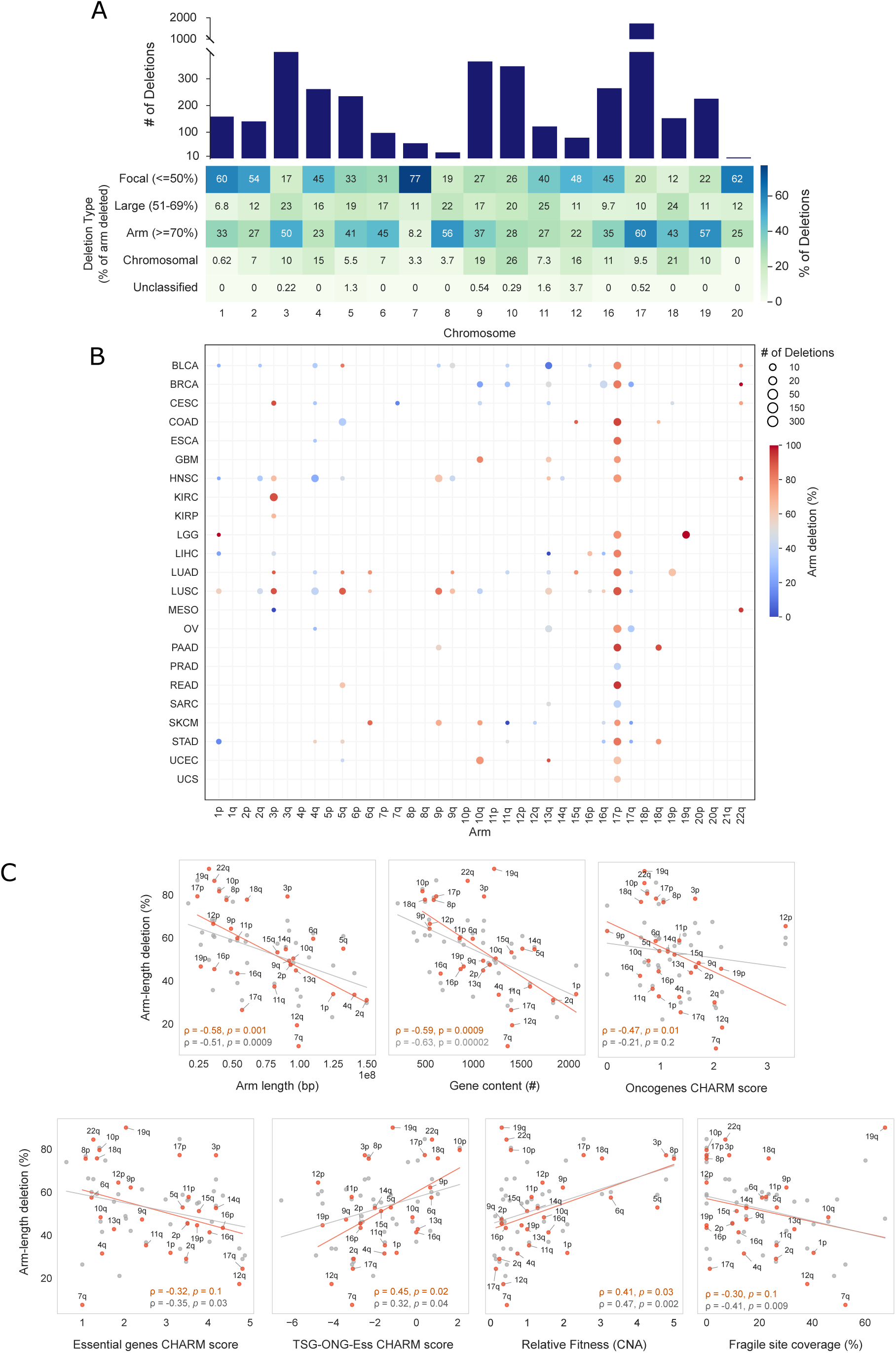
Chromosomal properties constrain deletion lengths in TSG two-hits. (A) Number of TSG two-hit events in each non-acrocentric chromosome (top), and the relative prevalence of associated deletion types stratified by the normalized deletion length or fraction of the arm deleted: focal (≤ 50%), large (51-69%), arm (≥ 70%), chromosomal (≥ 70% of both arms), unclassified (≤ 70% of both arms with ≤ 20% difference between arms) (bottom). (B) Prevalence of arm-length deletions in TSG two-hit events, defined as the percentage of deletions that are arm-length versus focal, shown separately for individual chromosome arms across cancer types. Bubble size reflects the total number of deletions (arm-length + focal). Only chromosome arms with ≥ 10 deletions are shown. (C) Relationships between pan-cancer arm-length deletion prevalence in TSGs (orange) and unclassified genes (grey), and chromosome arm–intrinsic properties, including arm length, gene content, CHARM scores, CNA-based net relative fitness values, and common fragile site coverage. Only chromosome arms with ≥ 10 deletions were included. Orange and gray lines represent linear regression fits for TSGs and unclassified genes, respectively. Insets show Spearman’s correlation coefficients and p-values.

**Fig. S6:**
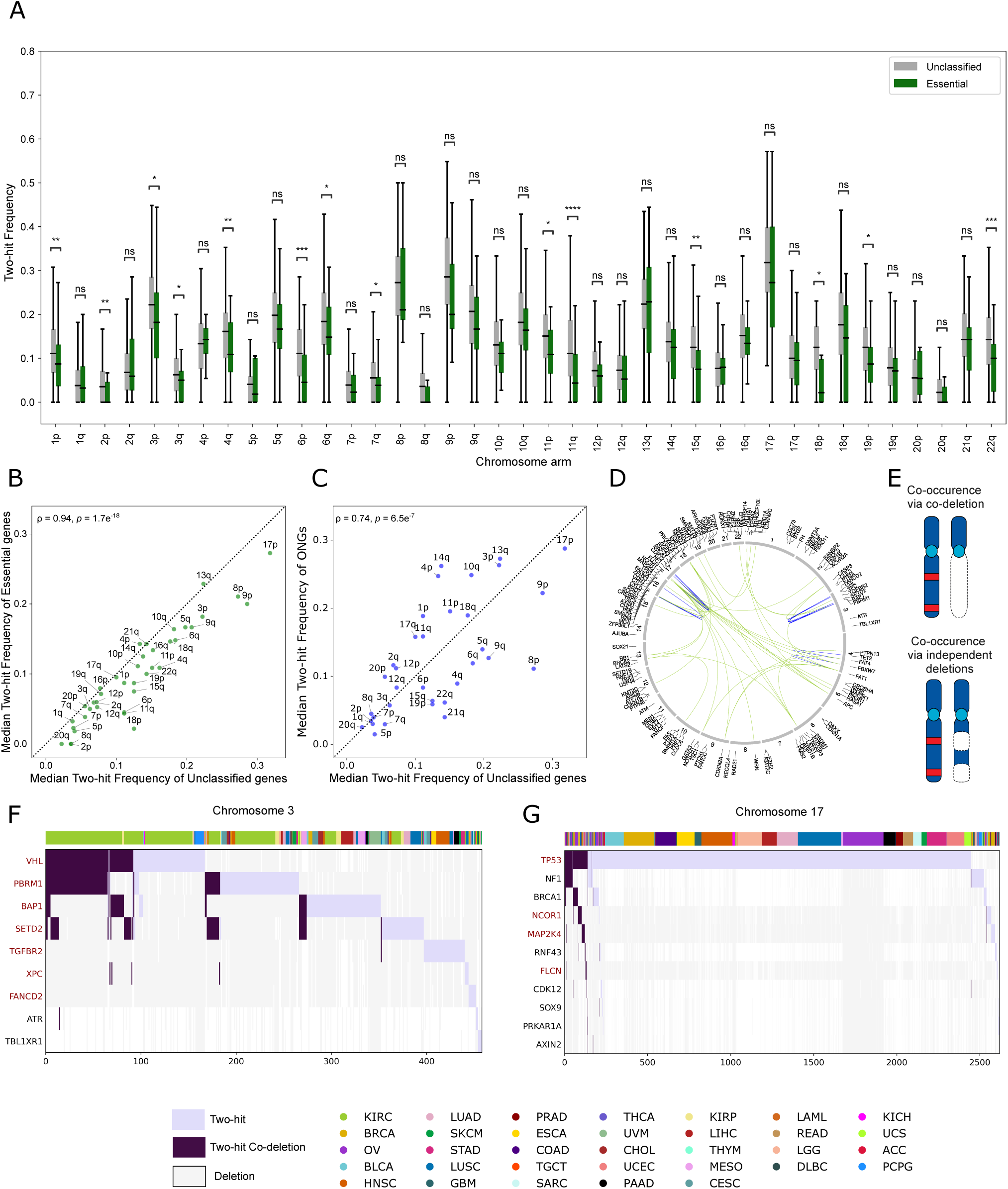
Chromosomal location influences TSG two-hit frequencies. (A) Pan-cancer distributions of two-hit frequencies for essential genes (green) and unclassified genes (grey) across chromosome arms. Asterisks denote statistical significance based on Mann–Whitney U tests: *p* > 0.05 (ns), ≤ 0.05 (*), ≤ 0.01 (**), ≤ 0.001 (***), ≤ 0.0001 (****). (B-C) Relationships between median pan-cancer two-hit frequencies of unclassified genes and those of essential genes (B) and oncogenes (C). Spearman’s correlation coefficients and p-values are shown in the insets. (D) Circos plot showing TSG pairs with significant co-occurrence of two-hit events (Fisher’s exact test, FDR < 0.05). Pairs located on the same chromosome are highlighted in blue. (E) Schematic illustrating two mechanisms of TSG two-hit co-occurrence: via a single co-deletion affecting multiple genes (top) or through independent deletions of each gene (bottom). Red bars denote point mutations, and dotted outlines denote deletions. (F-G) Oncoprint plots showing co-occurrence and co-deletion of TSG two-hit events on chromosomes 3 (F) and 17 (G). Each purple bar indicates a two-hit event for a given TSG in an individual patient, with darker shading representing co-deletions underlying co-occurring two-hit events. Grey bars represent deletions not accompanied by point mutations. Color bars at the top indicate cancer types. P-arm genes are highlighted in red.

**Fig. S7:**
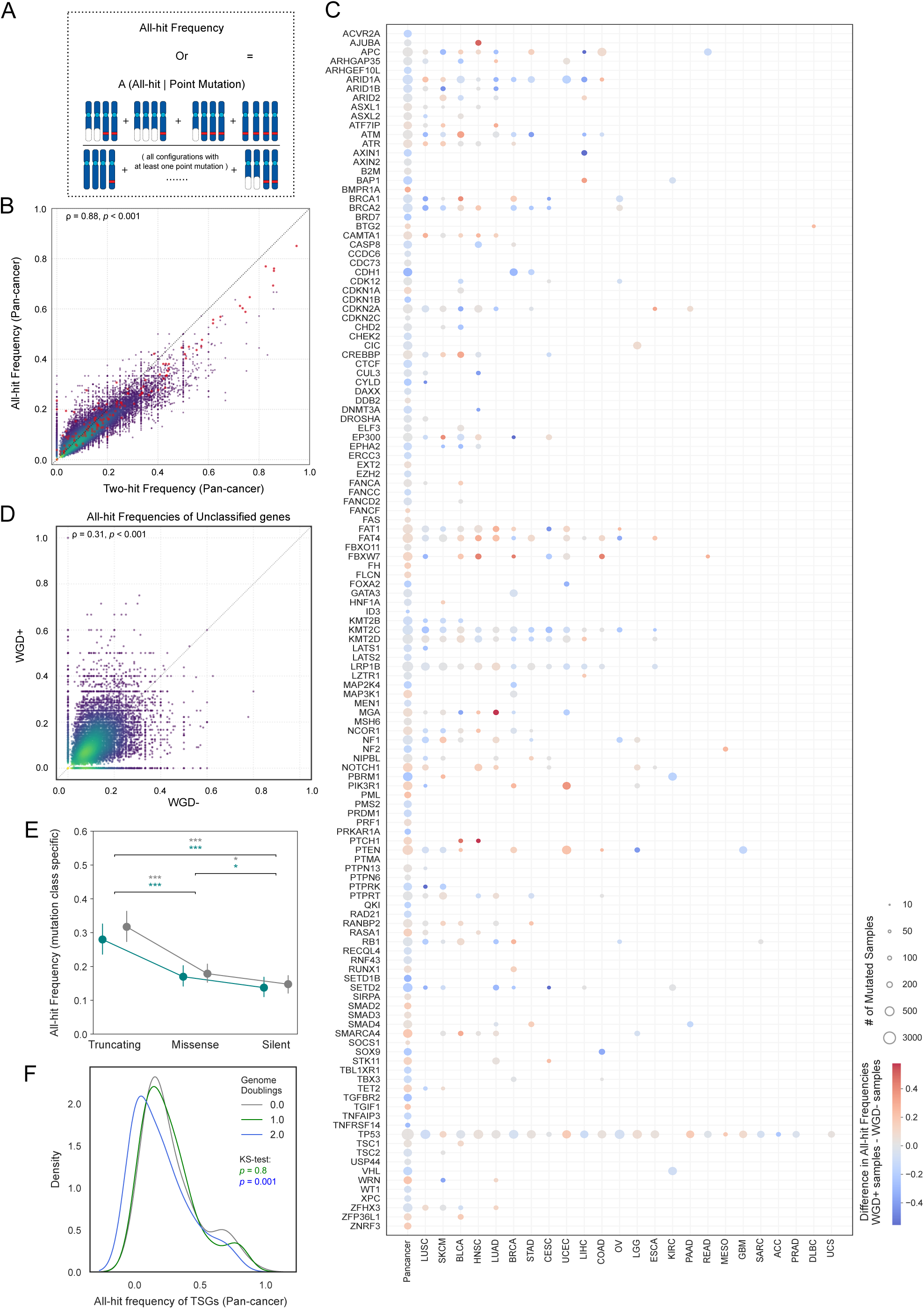
Whole genome doubling and all-hit frequencies. (A) Schematic illustrating the computation of all-hit frequencies for each gene at pan-cancer or cancer type levels. (B) Density scatter plot comparing pan-cancer two-hit and all-hit frequencies across all genes. Insets report Spearman’s correlation coefficients and corresponding p-values. TSGs are highlighted in red. (C) Bubble plot showing differences in all-hit frequencies between WGD+ and WGD- tumors across TSG–cancer-type pairs. Darker red indicates higher all-hit frequencies in WGD+ tumors, whereas darker blue indicates higher frequencies in WGD- tumors. Bubble size denotes the number of samples harboring point mutations. (D) Density scatter plot comparing pan-cancer all-hit frequencies of unclassified genes between WGD+ and WGD- tumors. All unclassified genes are shown. Insets report Spearman’s correlation coefficient and corresponding p-value. (E) Distributions of pan-cancer all-hit frequencies for TSGs, stratified by mutation class and WGD status. Points and bars denote the median and standard deviation, respectively. Asterisks indicate statistical significance from paired Wilcoxon signed-rank tests, with colors corresponding to WGD+ and WGD- groups: *p* > 0.05 (ns), ≤ 0.05 (*), ≤ 0.001 (***) (F) Kernel Density Estimation (KDE) plots showing TSG all-hit frequency distributions in samples stratified by the number of genome doubling events. Insets display Kolmogorov–Smirnov (KS) test p-values for comparisons between GD0 vs GD1 (green), and GD0 vs GD2 (blue).

**Fig. S8:**
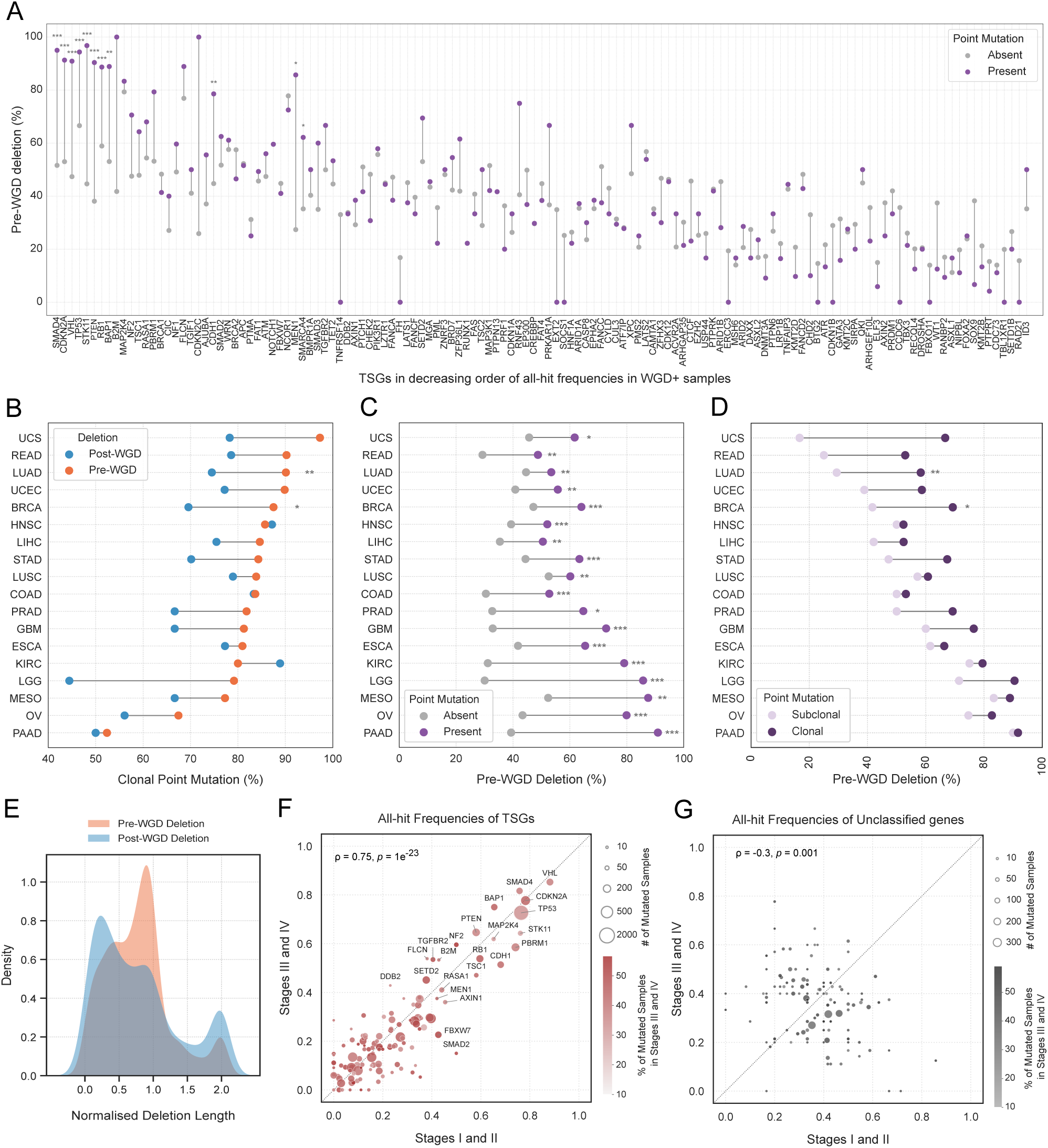
Temporal dynamics of alterations in whole genome doubled tumors. (A) Prevalence of pre-WGD deletions in the presence vs absence of point mutation in the remaining allele, shown across TSGs ordered by decreasing all-hit frequencies in WGD+ tumors. Asterisks indicate genes with significant differences between groups based on Fisher’s exact test with FDR correction (Benjamini-Hochberg method): *p* ≤ 0.05 (*), ≤ 0.01 (**), ≤ 0.001 (***). (B-D) Cancer-type–specific prevalence of: clonal point mutations co-occurring with pre-WGD vs post-WGD deletions of the remaining allele in TSGs (B); pre-WGD TSG deletions in the presence vs absence of point mutations in the remaining allele (C); and pre-WGD TSG deletions accompanying clonal vs subclonal point mutations in the remaining allele (D). Asterisks denote cancer types with significant differences between groups based on Fisher’s exact test with FDR correction (Benjamini-Hochberg method): *p* ≤ 0.05 (*), ≤ 0.01 (**), ≤ 0.001 (***). Only cancer types significant in at least one of the three comparisons are shown. (E) Kernel density estimation (KDE) plots showing normalized deletion length distributions for pre- and post-WGD deletions in TSG all-hit events across all non-acrocentric chromosomes. The area under each curve equals 1. (F-G) Pan-cancer all-hit frequencies of TSGs (F) and unclassified genes (G) in early-stage (I–II) vs late-stage (III–IV) tumors. Only unclassified genes with two-hit or all-hit frequencies ≥ 0.4 across all samples (irrespective of stage) are shown. Bubble size represents the number of samples with point mutations, and bubble hue indicates the percentage of mutated samples are late-stage (III–IV). Insets report Spearman’s correlation coefficients and corresponding *p*-values.

